# PI(4,5)P_2_ role in Transverse-tubule membrane formation and muscle function

**DOI:** 10.1101/2024.01.31.578124

**Authors:** Naonobu Fujita, Shravan Girada, Georg Vogler, Rolf Bodmer, Amy A. Kiger

## Abstract

Transverse (T)-tubules – vast, tubulated domains of the muscle plasma membrane – are critical to maintain healthy skeletal and heart contractions. How the intricate T-tubule membranes are formed is not well understood, with challenges to systematically interrogate in muscle. We established the use of intact Drosophila larval body wall muscles as an ideal system to discover mechanisms that sculpt and maintain the T-tubule membrane network. A muscle-targeted genetic screen identified specific phosphoinositide lipid regulators necessary for T-tubule organization and muscle function. We show that a *PI4KIIIα*-*Skittles/PIP5K* pathway is needed for T-tubule localized PI(4)P to PI(4,5)P_2_ synthesis, T-tubule organization, calcium regulation, and muscle and heart rate functions. Muscles deficient for *PI4KIIIα* or *Amphiphysin*, the homolog of human *BIN1*, similarly exhibited specific loss of transversal T-tubule membranes and dyad junctions, yet retained longitudinal membranes and the associated dyads. Our results highlight the power of live muscle studies, uncovering distinct mechanisms and functions for sub-compartments of the T-tubule network relevant to human myopathy.

**Summary:** T-tubules – vast, tubulated domains of the muscle plasma membrane – are critical to maintain skeletal and heart contractions. Fujita *et al*. establish genetic screens and assays in intact Drosophila muscles that uncover PI(4,5)P_2_ regulation critical for T-tubule maintenance and function.

**Key Findings:**

- *PI4KIIIα* is required for muscle T-tubule formation and larval mobility.
- A *PI4KIIIα-Sktl* pathway promotes PI(4)P and PI(4,5)P_2_ function at T-tubules.
- *PI4KIIIα* is necessary for calcium dynamics and transversal but not longitudinal dyads.
- Disruption of PI(4,5)P_2_ function in fly heart leads to fragmented T-tubules and abnormal heart rate.

## Introduction

Muscle function relies on specialized organelles. Two muscle-specific membrane compartments – the transverse (T)-tubules and sarcoplasmic reticulum (SR) – are essential to power contractions. T-tubules are tubulated domains of the muscle cell membrane that invaginate radially inward along the central sarcomeres. Cell surface signals relayed at T-tubule and SR membrane contact sites, called dyad or triad junctions, trigger the coordinated calcium (Ca^2+^) release that enables synchronous sarcomere contractions. Proper formation and organization of the T-tubule network is thus critical for both skeletal and heart muscle function (Dibb et al., 2022, Al-Qusairi and Laporte, 2011).

The T-tubule membrane network is vast, containing a large proportion of the total myofiber plasma membrane. The network includes transversal and longitudinal branched membrane elements that vary depending on different muscle stages and types. In mouse embryonic skeletal muscle, T-tubules first appear as longitudinal membranes that then remodel postnatally to include mainly transversal elements (Takekura et al., 2001). In contrast, both longitudinal and transversal T-tubule elements are maintained in adult mammalian cardiac muscle (Brette and Orchard, 2007) and insect muscles (Jensen, 1977, Takekura and Franzini-Armstrong, 2002). Interestingly, a reversion of the T-tubule network from more transversal to more longitudinal elements corresponded with certain pathological states in both cardiomyocytes (Lipsett et al., 2019, Wei et al., 2010) and skeletal muscle (Al-Qusairi et al., 2009, Chin et al., 2015, Dowling et al., 2009, Fugier et al., 2011). Altogether, the progression and reversion of T-tubule organization, respectively, raises the possibility that the transversal and longitudinal elements may serve distinct contributions in the membrane network.

The striking network organization and uniquely tubulated membranes suggest there are discrete mechanisms for T-tubule membrane biogenesis. Although continuous extensions of the plasma membrane, T-tubules exhibit an enrichment over the plasma membrane in cholesterol, the BAR domain protein, BIN1/Amphiphysin, and the L-type Ca^2+^ channel (Al-Qusairi and Laporte, 2011, Lee et al., 2002, Rosemblatt et al., 1981). How are T-tubule membranes specified and differentiated into these distinct domains? General determinants and organizers of plasma membrane identity are implicated. BIN1/AMPH2 has a conserved function in membrane tubulation and a broadly demonstrated requirement for T-tubule formation in heart and skeletal muscle Butler-(Lee et al., 2002, Muller et al., 2003, Razzaq et al., 2001). In addition, there are proposed T-tubule roles for other membrane-related protein functions of MTM1, CAV3, DYSF, EHD and DNM2 (Al-Qusairi et al., 2009, Chin et al., 2015, Galbiati et al., 2001, Hofhuis et al., 2017, Klinge et al., 2010, Posey et al., 2014, Ribeiro et al., 2011). Mutations in the human homologs of each also are associated with myopathy and/or cardiomyopathy (Bashir et al., 1998, Betz et al., 2001, Bitoun et al., 2005, Laporte et al., 1996, Liu et al., 1998, Minetti et al., 1998, Nicot et al., 2007), pointing to the importance of membrane regulation for muscle organization and function.

Combining genetics with microscopy imaging in intact muscle is a powerful approach for discovery of T-tubule functions, as recently demonstrated by the tractable *in vivo* screens in fruit flies and zebrafish (Fujita et al., 2017, Hall et al., 2020). Drosophila has proven insightful for uncovering requirements specific for T-tubule formation and T-tubule remodeling. First, gene functions required for T-tubule membrane formation are conserved. The conserved role for BIN1/AMPH2 in T-tubule formation was first described for the single Drosophila homolog, Amphiphysin (Amph) (Razzaq et al., 2001). The *Amph* null mutant flies lack transversal T-tubule membranes in myofibers at all developmental stages, corresponding with defects in larval mobility and adult flight (Razzaq et al., 2001). We showed that T-tubules undergo a regulated remodeling in a subset of larval body wall muscles during metamorphosis, and that *Amph* is required again to reform T-tubules with remodeling (Fujita et al., 2017, Ribeiro et al., 2011). In contrast to *Amph*, we found Drosophila *mtm*, a phosphoinositide 3-phosphatase, is not involved in initial T-tubule biogenesis, but is only required for T-tubule membrane reformation with muscle remodeling (Ribeiro et al., 2011). Second, T-tubules in intact Drosophila muscles are easily imaged directly through the cuticle of live animals. Using genetic screens by microscopy imaging in intact muscle, we also identified additional new functions needed for T-tubule membrane network organization (Fujita et al., 2017). Our studies show that T-tubule formation and remodeling involve overlapping yet distinct requirements, such as seen for *Amph* versus *mtm* functions, respectively, and that genetic programs for formation versus remodeling can be studied at different developmental stages in Drosophila body wall muscle.

Here, we screened phosphoinositide lipid regulators for roles in T-tubule membrane organization in intact Drosophila muscle. Our results identified that T-tubule-localized phosphoinositide synthesis of PI(4)P to PI(4,5)P_2_ is required for transversal tubulated domains, dyad formation, calcium flux, muscle contraction and animal mobility. Our results underscore the importance of membrane lipid identity in muscle function. Thus, Drosophila provides unique advantages for genetic and imaging approaches in differentiated muscle to uncover mechanisms for T-tubule membrane organization that impact muscle function.

## Results and Discussion

### Screen identifies *PI4KIIIα* required for T-tubule membranes and larval mobility

Drosophila larval body wall muscles are individual myofibers with a well-formed T-tubule network. The body wall muscles span each segment just beneath the transparent larval cuticle, permitting an advantageous combination of muscle-targeted expression (Mef2-GAL4) and microscopy imaging in intact animals. To test the genetic basis for T-tubule membrane organization, we established use of the dorsal internal oblique muscles (hereafter, simply referred to as larval body wall muscles or just muscles). When imaged in live, third instar larval muscles, the Dlg1:GFP endogenous-trap fusion protein was detectable along the characteristic T-tubule membrane network with tubular elements spanning both transversal and longitudinal directions (**Figure 1A; Supplemental video 1**). Dlg1:GFP localized with Amphiphysin (Amph) at T-tubule membranes that regularly traversed the array of sarcomeres (Zormin/z-line; **Figure 1B-1C**). Muscle-targeted *Amph* knockdown led to loss of the Dlg1:GFP-marked transversal tubules (**Figure 1D**), as predicted (Razzaq et al., 2001), indicating the ability to identify potent RNAi-induced T-tubule defects by live muscle imaging.

**Figure 1.**
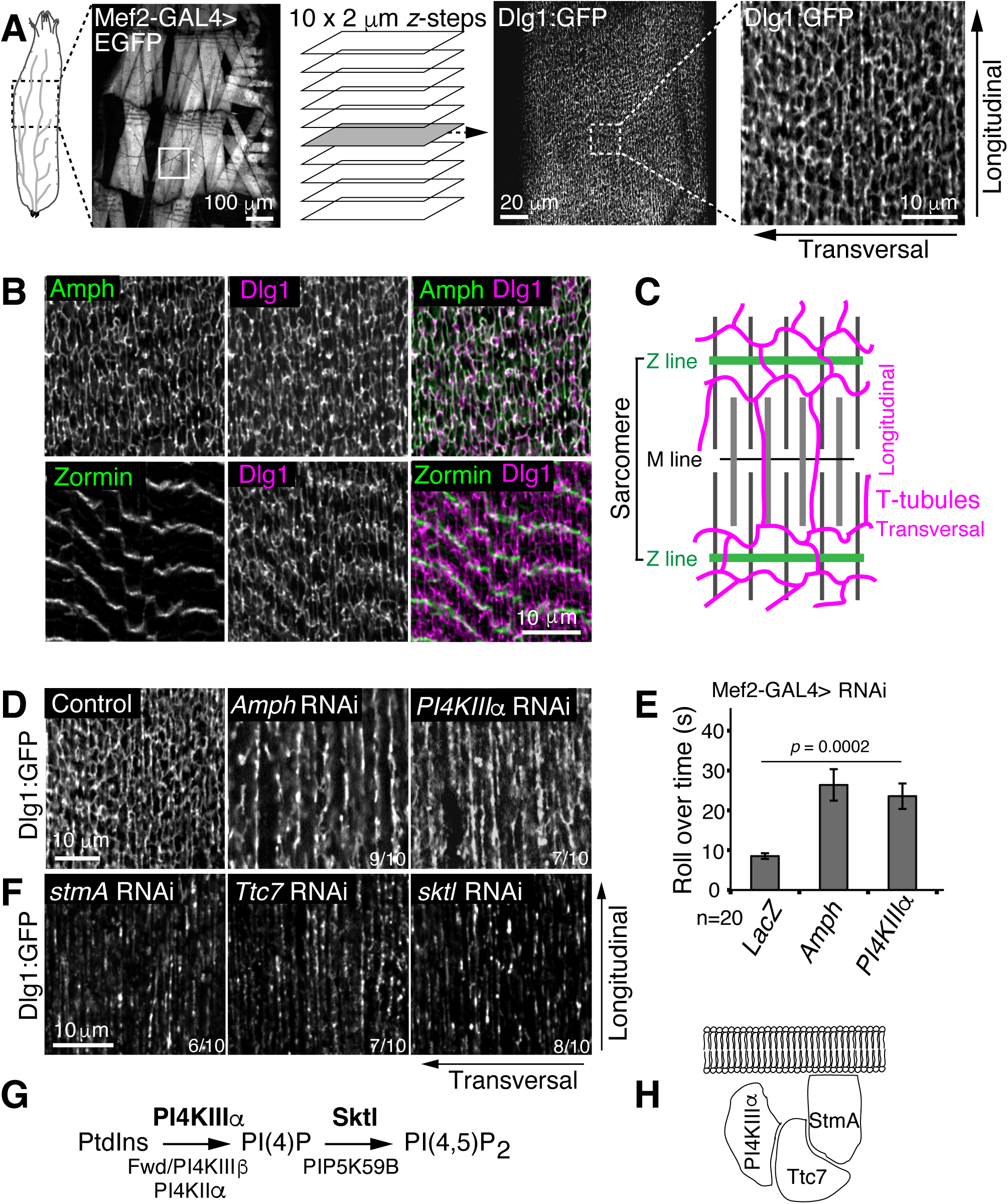
*PI4KIIIα-sktl* pathway required for T-tubule membranes and larval mobility. **(A)** RNAi screen in larval body wall muscles, using Mef2-GAL4 (left, UAS-EGFP) and confocal imaging of Dlg1:GFP detection of T-tubules (right). Live, intact dorsal longitudinal muscles in abdominal segments imaged as z-stacks through the larval cuticle, as shown in **Supplemental video 1**. Arrows indicate orientation of transversal and longitudinal T-tubule membrane elements. **(B)** Dlg1 (pink) and Amph (green) immunostaining co-localized at T-tubule membranes (top) throughout sarcomeres (bottom; anti-Zormin, green). **(C)** Cartoon of T-tubule network organization (pink), with regular intervals of transversal membranes adjacent to the sarcomere Z-lines (green) and longitudinal membranes between myofibrils. **(D)** Dlg1:GFP primary screen images. Control: *DMef2-GAL4, UAS-lacZ*. RNAi of *Amph* and *PI4KIIIα* both showed loss of transversal membranes and a thickening of longitudinal membranes. Numbers indicate images with observed defects from total images screened. **(E)** Mean larval roll over time (seconds). Individual larvae placed dorsal-side down and timed how long to roll over to dorsal-side up. SD from 3 experiments. *p* value = 0.0002. **(F)** Dlg1:GFP with *stmA* RNAi, *Ttc7* RNAi or *sktl* RNAi. **(G)** Pathway for conversion of PtdIns to PI(4)P then PI(4,5)P_2_ by possible Drosophila PI4-kinases and PIP5-kinases, respectively; muscle RNAi of only *PI4KIIIα* and *sktl* resulted in disrupted T-tubules. **(H)** Drosophila PI4KIIIα scaffold proteins, StmA and Ttc7 (also known as Efr3/RBO and YPP1, respectively). Scale bars 10 µm, except as indicated 100 µm and 20 µm in (A).

To discover genes involved in forming and maintaining T-tubules, we screened RNAi lines targeting predicted membrane-associated functions. Given the relevance of phosphoinositide (PIP) lipids and their regulators to membrane identity (Balla, 2013, Hammond and Burke, 2020) and muscle health (Blondeau et al., 2000, Kwok et al., 2023, Laporte et al., 2003, Ribeiro et al., 2011, Szentesi et al., 2023, Taylor et al., 2000, Voelker et al., 2023), we focused on effects from knockdown of 28 Drosophila phosphoinositide kinase and phosphatase family members. From 52 RNAi lines collectively targeting 27 Drosophila PIP regulators, we were able to image and score 47 RNAi conditions in third instar larval muscle for 26 candidate genes (**Table 1**).

**Table 1.** RNAi survey of T-tubule organization in third instar larval muscle (Mef2-GAL4). List of Drosophila phosphoinositide kinases and phosphatases by gene name, CG-number and predicted closest human ortholog by percent identity in amino acid sequence and enzymatic activity. Tested RNAi stocks listed by genotype and stock center number from Bloomington Drosophila Stock Center (BDSC) or Vienna Drosophila Resource Center (VDRC), or as shown. Phenotypes scored for animal viability or lethal stage, and for observed Dlg1:GFP marked T-tubule organization as imaged through the cuticle in live intact muscle. ND = not determined, either due to lethality or unavailable stock. Table information based on genotypes in FlyBase (FB2023_04).

Among the lines surveyed, knockdown of *PI4KIIIα* resulted in muscles lacking transversal T-tubules and a thickening of the longitudinal membranes, a strikingly similar phenotype to that seen in *Amph-*depleted muscle (**Figure 1D, Supplemental Figure S1A**). Likewise, muscle-targeted *Amph* and *PI4KIIIα* RNAi both similarly resulted in impaired muscle function with an increased larval roll-over time (**Figure 1E**), consistent with known mobility defects of *Amph* null mutants (Razzaq et al., 2001). *PI4KIIIα* knockdown examined at earlier larval stages also showed fewer transversal elements and their loss with larval muscle growth (**Supplemental Figure S1B**). These results implied shared requirements for *PI4KIIIα* and *Amph* in T-tubule membrane formation and/or maintenance.

### *PI4KIIIα-sktl* pathway promotes PI(4,5)P_2_ function for T-tubule membrane formation

PI4KIIIα is one of three Drosophila PI4-kinases able to phosphorylate phosphatidylinositol to phosphatidylinositol 4-phosphate, or PI(4)P. We validated the screen result with multiple RNAi lines against the three genes encoding PI4-kinases, *PI4KIIIα*, *fwd/PI4KIIIβ* or *PI4KIIα*, demonstrating a *PI4KIIIα*-specific requirement for T-tubule membranes (**Supplemental Figure S1C**). PI4-kinase synthesis of PI(4)P could have direct functions and/or serve as a PIP5-kinase substrate for synthesis of PI(4,5)P_2_. To distinguish how *PI4KIIIα* function is needed for T-tubule organization, we disrupted members of a PI4KIIIα pathway involved in PI(4,5)P_2_ functions at the plasma membrane (Baird et al., 2008, Balakrishnan et al., 2018, Devergne et al., 2014, Liu et al., 2018, Nakatsu et al., 2012, Tan et al., 2014, Wu et al., 2014b, Yan et al., 2011, Zhang et al., 2017). First, we found a shared loss of transversal T-tubule membranes with knockdown of known PI4KIIIα plasma membrane scaffold proteins, StmA and TTC7 (homologs also known as Efr3/RBO and YPP1, respectively (Huang et al., 2004, Liu et al., 2018)) (**Figure 1F-1H**). Second, specific depletion of the *skittles* (*sktl*) PIP5-kinase, but not *PIP5K59*, also led to a lack of transversal T-tubules (**Figure 1F-1G**; **Supplemental Figure S1C**). In each case, muscles with T-tubule knockdown phenotypes still contained striated F-actin myofibrils, indicating that overall muscle development and organization were largely unaffected (**Supplemental Figure S1D**). Altogether, dependence on the *PI4KIIIa-sktl* pathway in larval muscle suggested an important PI(4,5)P_2_ role for T-tubule membrane organization.

### PI(4)P and PI(4,5)P_2_ at T-tubules depends on *PI4KIIIa-sktl* pathway

Phosphoinositide lipids – the seven different phosphorylated forms of phosphatidylinositol – are differentially enriched at membrane domains that direct localized functions (Balla, 2013, Hammond and Burke, 2020). Since PIPs are interconverted by the selective and overlapping activities of PI-kinase and PI-phosphatase family members, the disruption of individual regulators can impact just specific pool(s) of distinct substrate(s) and/or product(s) (**Supplemental Figure S2A**).

In conjunction with the RNAi screen for functions needed for T-tubule organization, we explored the distribution of the different PIP species in wildtype muscle. We surveyed the myofiber localization of eight selective biosensors available for five of the seven different PIPs, including PI(3)P, PI(4)P, PI(3,4)P_2_, PI(4,5)P_2_ and PI(3,4,5)P_3_. Only the PI(4)P indicator, P4M (Brombacher et al., 2009, Hammond et al., 2014, Liu et al., 2018), and the two PI(4,5)P_2_ indicators, GFP-tagged PH-PLCδ and Tubby-C (Santagata et al., 2001, Stauffer et al., 1998, Thallmair et al., 2022, Varnai and Balla, 1998), were found to localize at the T-tubule network in larval body wall muscles (**Supplemental Figure S2B**). In wildtype conditions, both PI(4)P and PI(4,5)P_2_ co-localized with Dlg1, as well as with each other, along T-tubule membranes (**Figure 2A-2E**).

**Figure 2.**
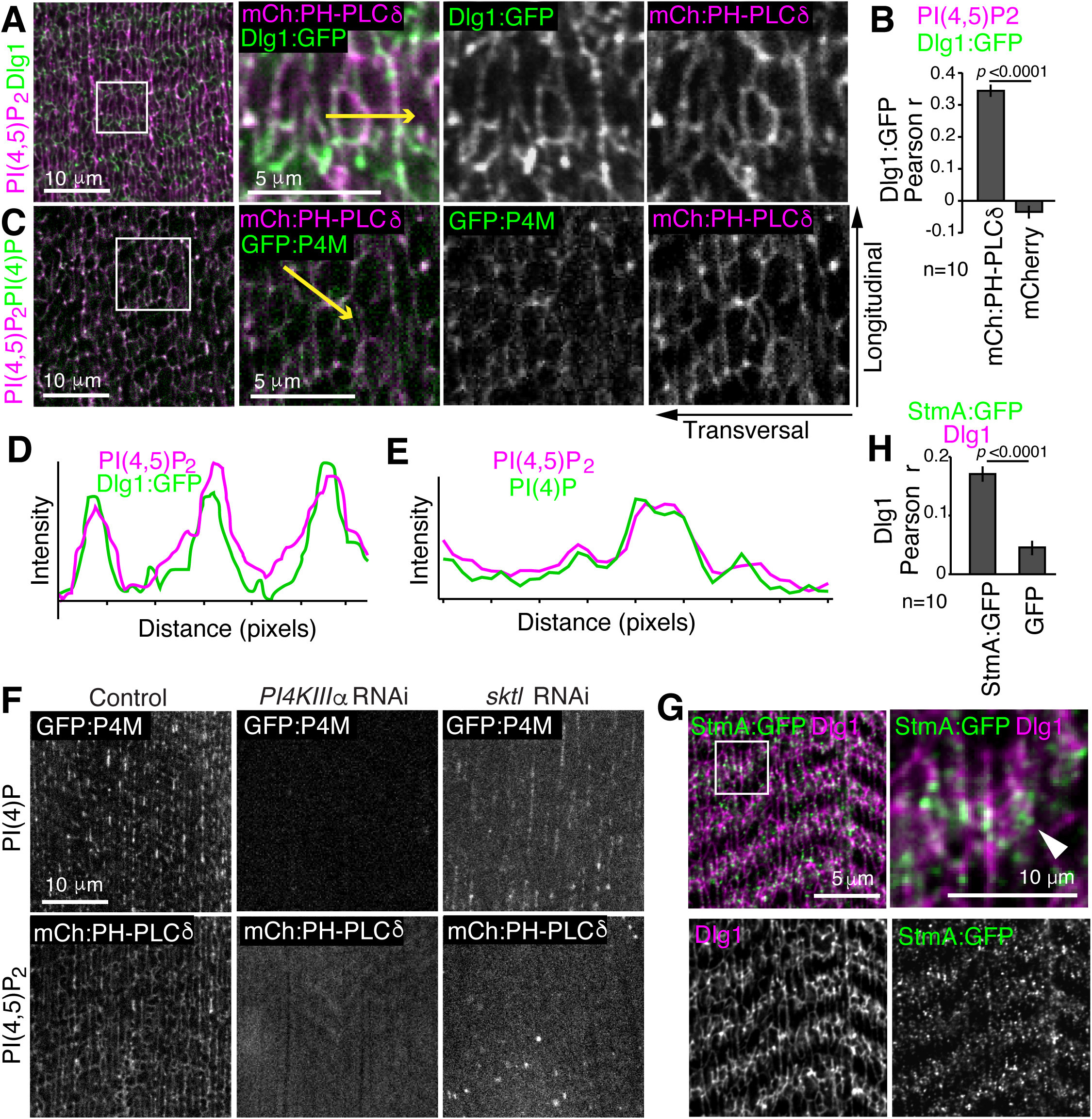
PI(4)P and PI(4,5)P_2_ accumulate at T-tubule membranes and depend on PI4KIIIα- Sktl pathway. **(A)** PI(4,5)P_2_ (mCherry:PH-PLC8, pink) is enriched along T-tubules (Dlg1:GFP, green); white-framed region (left) shown as crops for merged and single channels (right). **(B)** Pearson correlation (*r*) between Dlg1:GFP and mCherry:PH-PLC8 or mCherry control. *p* value < 0.0001. **(C)** PI(4,5)P_2_ (mCherry:PH-PLC8, pink) colocalizes with PI(4)P (GFP:P4M, green); white framed region (left) shown as crops for merged and single channels (right). **(D-E)** Line tracings of pixel intensities along yellow arrows drawn in (A) and (C), respectively. **(F)** PI(4)P (GFP:P4M, top row) is at T-tubules in control muscle (*lacZ*), absent in *PI4KIIIα* RNAi muscle, but still present in *sktl* RNAi muscle. PI(4,5)P_2_ (mCherry:PH-PLC8, bottom row) is at T-tubules in control muscle (*lacZ*), but absent in both *PI4KIIIα* RNAi and *sktl* RNAi muscles. **(G)** StmA:GFP (green) forms clusters of foci along T-tubules (anti-Dlg1, pink) in merge (top, arrowhead) and single channel images (below); white framed region (left) shown as crop (right). **(H)** Pearson correlation (*r*) between anti-Dlg1 and StmA:GFP or GFP control. *p* value < 0.0001. Scale bars 10 µm, except as indicated 5 µm for (G) and crops in (A) and (C).

Muscles with disruption of the *PI4KIIIα*-*sktl* pathway members or *Amph* RNAi exhibited loss of the Dlg1-marked transversal pattern yet retained abnormally thick longitudinal T-tubule membranes (**Figure 1F**). We tested whether this loss of the Dlg1-marked transversal elements corresponded with changes to the normal PI(4)P and/or PI(4,5)P_2_ T-tubule network pattern. While *PI4KIIIα* and *stmA* RNAi conditions led to the complete loss of PI(4)P at both transversal and longitudinal elements, the *sktl* RNAi condition showed loss of just the transversal PI(4)P pattern but retained the longitudinal pattern (**Figure 2F**, top). In contrast, all three conditions – *PI4KIIIα*, *stmA* and *sktl* RNAi – each resulted in severe depletion of the PI(4,5)P_2_ pattern throughout the entire T-tubule network (**Figure 2F, Supplemental Figure S2C**). Thus, the common features underlying disruption of the *PI4KIIIα*-*sktl* pathway was lack of T-tubule-associated PI(4,5)P_2_ and loss of the transversal membranes, supporting a key function for PI(4,5)P_2_ in T-tubule organization.

PI(4,5)P_2_ plays broad roles at the plasma membrane (Hammond and Burke, 2020, Katan and Cockcroft, 2020). Amph protein homolog, AMPH2/BIN1, can bind PI(4,5)P_2_ and was shown dependent on the phosphoinositide binding module to localize at T-tubules (Lee et al., 2002). We found that *Amph*-depleted muscle retained PH-PLCδ/PI(4,5)P_2_-marked longitudinal elements (**Supplemental Figure S2C**), supporting that *Amph* could act downstream from or in parallel to the *PI4KIIIα*-*sktl* pathway. Interestingly in all cases, muscles lacking transversal T-tubule membranes still retained Dlg1-marked longitudinal membranes. These results revealed there are distinct identities and regulation for longitudinal versus transversal T-tubule membrane elements.

To explore the distribution of PI4KIIIα activity in myofibers, we examined localization of the PI4KIIIα scaffold protein, StmA. In wildtype myofibers, StmA:GFP was detected at discrete clusters of foci along Dlg1-marked T-tubules (**Figure 2G-2H)**. This suggested that localized PI(4)P-synthesis at T-tubules underlies the PI(4,5)P_2_ function needed for transversal T-tubule network organization and function.

### *PI4KIIIα* is required for transversal but not longitudinal dyad junctions

Given both the distinct *PI4KIIIα* requirement for transversal membranes and the localization of StmA at T-tubules, we considered what effects *PI4KIIIα* function has on excitation-contraction coupling. The T-tubule and sarcoplasmic reticulum (SR) membrane contact sites called dyad junctions, analogous to the vertebrate triad junctions, mediate local calcium dynamics for excitation-contraction coupling along sarcomeres (Peachey, 1968, Smith, 1966). As an indicator of the dyads, we detected Junctophilin (Jp), a conserved SR-anchored junction protein (Calpena et al., 2018, Piggott and Jin, 2021, Takeshima et al., 2000). In wildtype muscle, Jp localized as expected to uniformly-distributed foci associated with both SR and T-tubule membranes (marked by Rtnl1 and Dlg1, respectively; **Figure 3A-3B; Supplemental Figure S3A**). Strikingly, Jp and StmA foci extensively colocalized with each other in wildtype muscle **(Figure 3C-3D**), suggesting that PI4KIIIα is recruited to T-tubules at sites of the dyad junctions.

**Figure 3.**
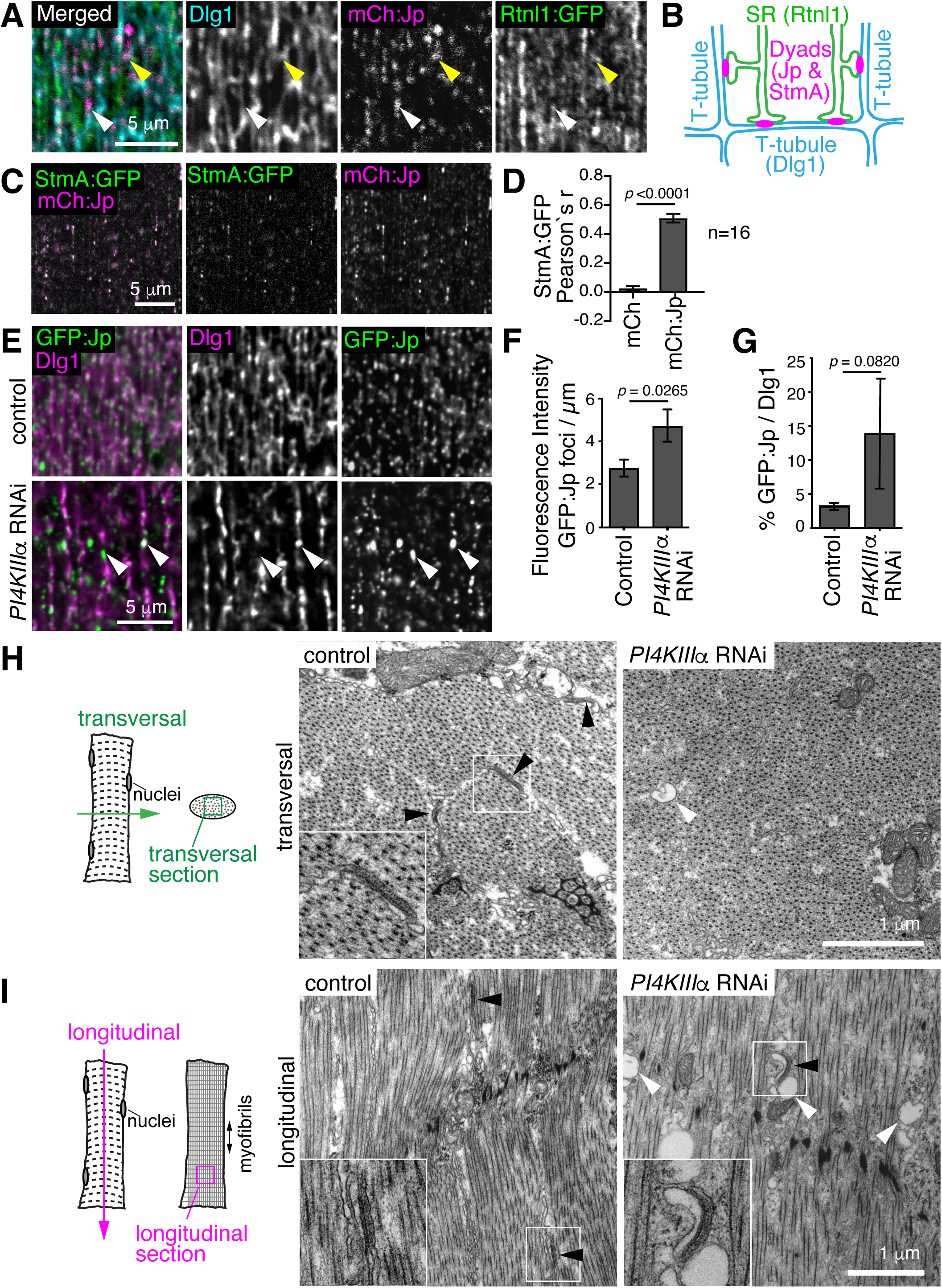
PI4KIIIα scaffolding at dyads is required for transversal membranes and normal dyad distribution. **(A)** Merge and single channels showing Junctophilin (mCherry:Jp, pink) at dyad junctions between the sarcoplasmic reticulum (SR; Rtnl1:GFP, green) and T-tubule membranes (anti-Dlg1, cyan). Jp associated with both transversal (yellow arrowheads) and longitudinal (white arrowheads) T-tubule membrane elements. **(B)** Cartoon of T-tubule and SR membrane contact sites at dyad junctions marked by Jp foci, where StmA was found colocalized. **(C)** StmA:GFP scaffold (green) colocalized with Jp-positive dyads (mCh:Jp, pink). **(D)** Pearson correlation (*r*) between StmA:GFP and control mCherry or mCherry:Jp. *p* value < 0.0001. **(E)** Dyads (GFP:Jp, green) and T-tubules (anti-Dlg1, pink) in control muscle (*lacZ*), showing GFP:Jp punctae of uniform size, distribution and intensity. In contrast, *PI4KIIIα* RNAi led to fewer, variably larger and more intense GFP:Jp regions along disrupted longitudinal T-tubules (arrowheads). **(F-G)** *PI4KIIIα* RNAi altered dyad distribution, with an increase in both the GFP:Jp foci fluorescence intensity per area (F) and the percentage colocalized with Dlg1 (G). **(H-I)** Transmission electron microscopy sections of control and *PI4KIIIα* RNAi larval body wall muscle. Black arrowheads and insets: electron-dense dyads. White arrowheads: swollen electron-light tubules. (H) Transversal sections in control show radial spokes of transversal T-tubule membranes and electron-dense dyads, both missing in *PI4KIIIα* RNAi. (I) Longitudinal sections contain dyads present in both control and *PI4KIIIα* RNAi muscle. Scale bars 5 µm (A, C, E) and 1 µm (H-I).

In *PI4KIIIα-*depleted myofibers, Jp foci were present but showed both an abnormal distribution pattern and morphology. The Jp foci were restricted to the longitudinal membranes and absent from the intervening regions, presumably due to loss of the transversal T-tubule membranes with *PI4KIIIα* depletion (**Figure 3E**). Compared to Jp foci in wildtype, the Jp foci in *PI4KIIIα* RNAi muscle were also enlarged with an expanded association at the longitudinal remnants (**Figure 3E-3G**). In contrast, the SR remained evenly distributed throughout *PI4KIIIα-*and *Amph-*depleted myofibers (**Supplemental Figure S3B**), suggesting specific effects on the T-tubule membranes and dyads.

Ultrastructural analysis in transversal sections of *PI4KIIIα-*depleted myofibers confirmed the near absence of electron-lucent, tubular transversal T-tubule membranes and any associated electron-dense dyad junctions (**Figure 3H**). In longitudinal sections, however, vacuolated electron-lucent membranes accumulated. Despite the abnormal morphology, electron-dense dyad junctions were seen along swollen longitudinal T-tubule membranes (**Figure 3I**). This confirms that in the absence of *PI4KIIIα*, dyads still formed at longitudinal T-tubule membranes despite a lack of PI(4,5)P_2_ and an abnormal membrane morphology.

### *PI4KIIIα* is necessary for calcium dynamics in excitation-contraction coupling

With loss of the transversal but not the longitudinal muscle dyad junctions, we wondered if calcium dynamics would be affected in *PI4KIIIα* knockdown muscle. Muscle contractions occur with cycles of increased cytoplasmic calcium, which is detectable by GFP fluorescence upon the GCaMP reporter binding (Nakai et al., 2001, Tian et al., 2009). Immobilized wildtype third instar larvae exhibited spontaneous muscle contractions coincident with peaks in Ca^2+^-dependent GCaMP6s signal (Chen et al., 2013) (**Figure 4A-4C; Supplemental video 2**). In contrast, larval body wall muscles with *Amph* RNAi or *PI4KIIIα* knockdown exhibited weaker contractions without such increases in cytoplasmic calcium (**Figure 4A-4C, Supplemental video 3**), consistent with the impaired larval mobility (**Figure 1E**). Similarly, *stmA* RNAi also disrupted calcium intracellular dynamics (**Figure 4C**), supporting the importance of PI4KIIIα activity at the T-tubule membrane for excitation-contraction coupling. Therefore, the presence of dyad junctions at abnormal longitudinal T-tubule membranes (**Figure 3**) are insufficient to support body wall muscle calcium dynamics, contractions or mobility functions.

**Figure 4.**
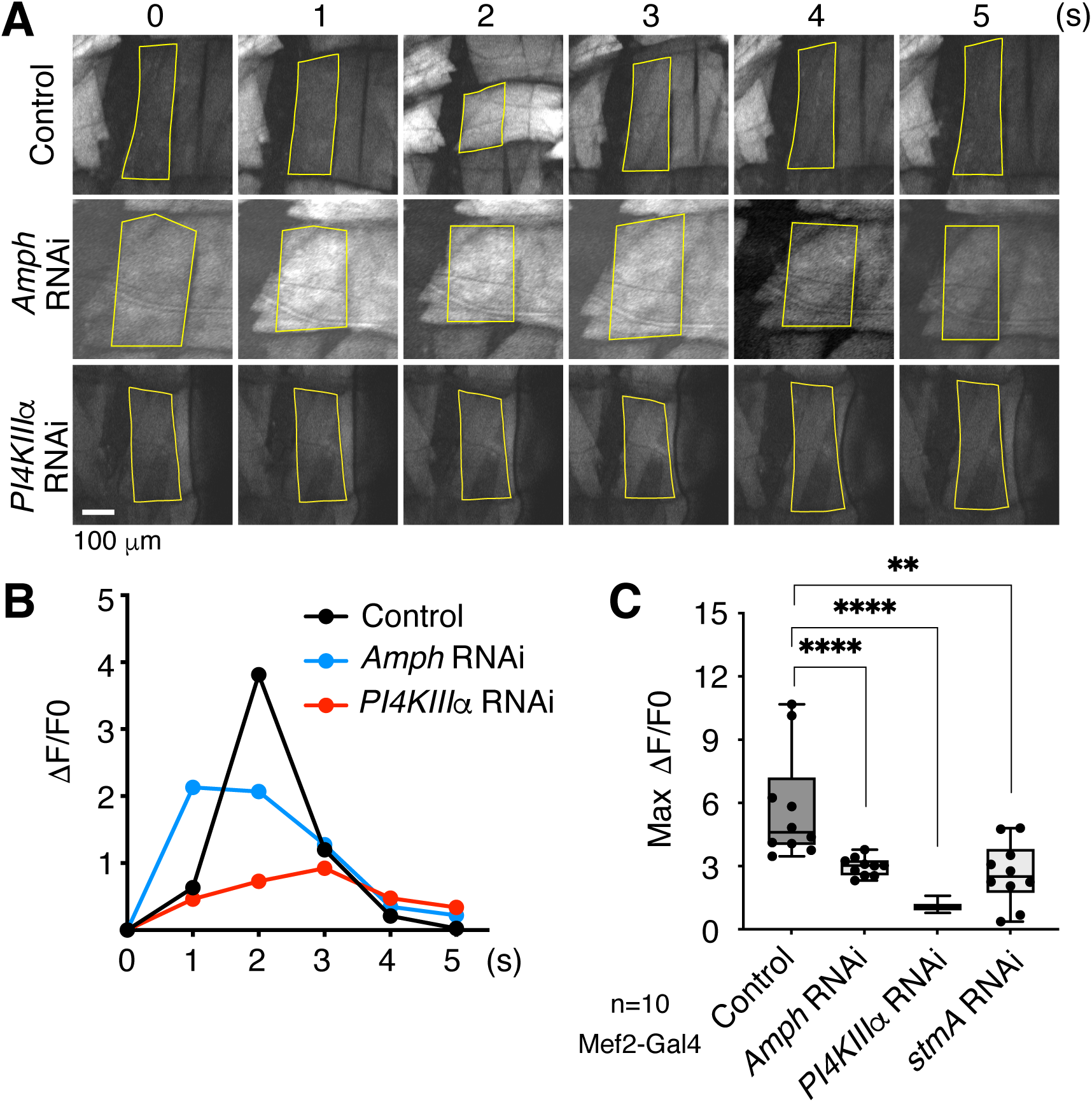
*PI4KIIIα* pathway is required for calcium dynamics with contraction. Mef2-GAL4 used for muscle-targeted RNAi and expression of GCaMP6s Ca^2+^-sensor. **(A)** Ca^2+^ dynamics in intact *lacZ* control, *Amph* RNAi, and *PI4KIIIα* RNAi dorsal larval muscle, shown as 1 sec frames from videos. Yellow lines, muscle regions used for quantification. Representative images taken from frames in **Supplemental video 2** and **Supplemental video 3**. **(B)** Change in GCaMP6s fluorescence intensity (ΔF) normalized to fluorescence intensity at time 0 (F0) shown over 1 sec video intervals within indicated muscle regions (shown in A) for *lacZ* control (black), *Amph* RNAi (blue) and *PI4KIIIα* RNAi (red). **(C)** Maximum change in GCaMP6s fluorescence intensity with spontaneous contractions imaged in *lacZ*, *Amph* RNAi, *PI4KIIIα* RNAi and *stmA* RNAi intact muscles (n=10 larvae). Box and whiskers with mean. ΔF/F0 = Fmax-Fmin/Fmin, ±SEM. Mann-Whiney tests for *p* values = 0.0084 (**), 0.0003 (***) and 0.0055 (**).

### PI(4,5)P_2_ depletion in adult heart leads to T-tubule disruption and cardiomyopathy

T-tubules are critical membrane domains for regulation of both skeletal and heart muscle function, and disruption of T-tubule organization is associated with forms of myopathy and cardiomyopathy (Al-Qusairi et al., 2009, Dibb et al., 2022). Drosophila have a simple heart, the dorsal vessel, in the shape of a tube made from pairs of cardiomyocytes (Souidi and Jagla, 2021). Larval and adult cardiomyocytes contain an elaborate T-tubule network seen by ultrastructure (Jensen, 1977, Lehmacher et al., 2012) and by the continuity of Dlg1-marked membranes (**Figure 5A**). In contrast, the cardiomyocyte T-tubule networks were fragmented and primarily longitudinal in dorsal vessels from *Amph* mutants or from animals with dorsal vessel-targeted *Amph* or *PI4KIIIα* RNAi (**Figure 5A; Supplemental Figure S3C**). High-speed imaging of the dorsal vessel (Fink et al., 2009) revealed that although spontaneous cardiac contractions still occurred (**Figure 5B; Supplemental Figure S4A-S4F**), there were significantly lower heart rates in *Amph* mutant hearts at one week and with heart-specific *Amph* RNAi at three and five weeks of age, respectively, indicative of cardiomyopathy (**Figure 5B-5C**).

**Figure 5.**
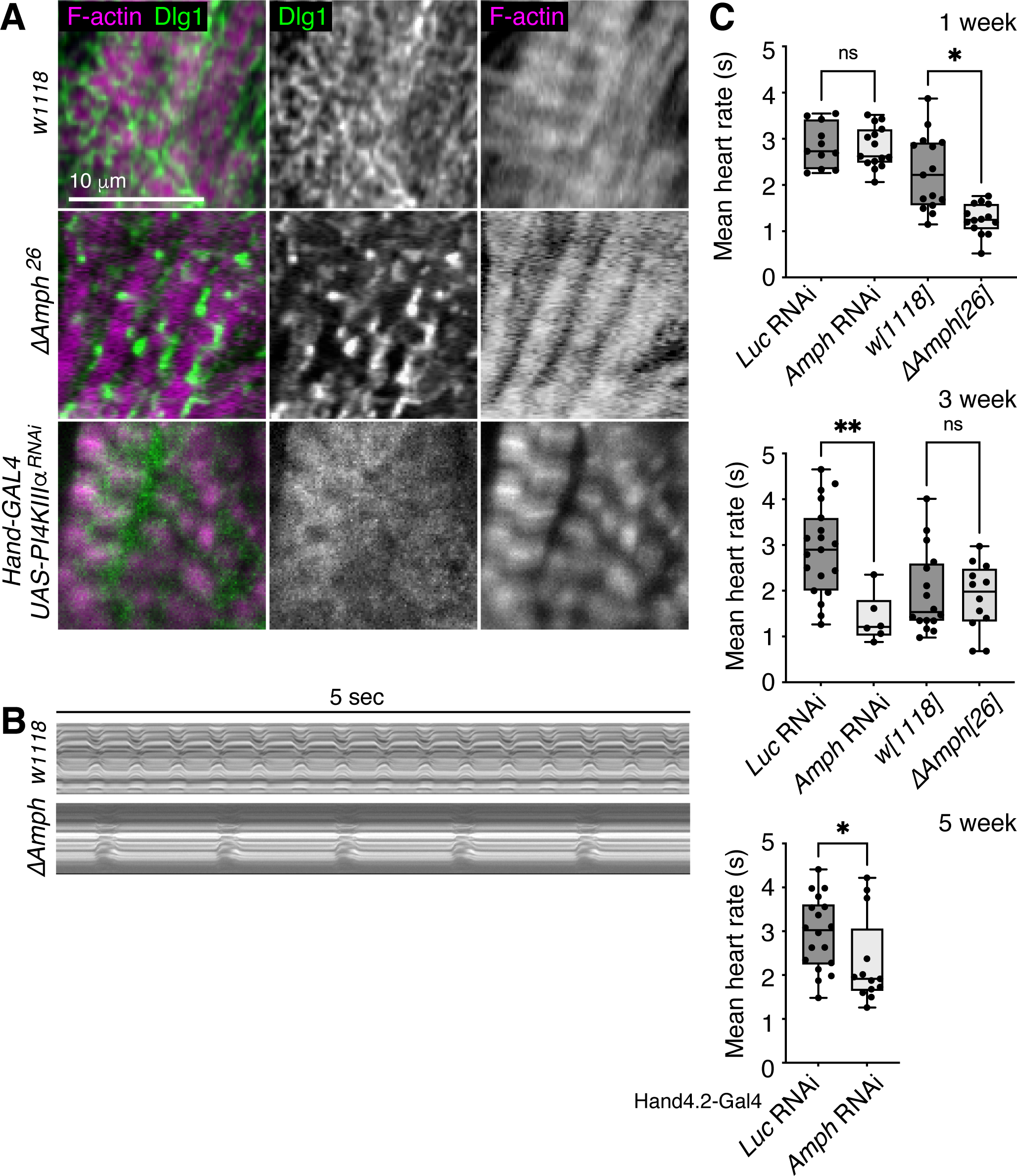
T-tubule PI(4,5)P_2_ roles in muscle predict requirements in heart function. **(A)** T-tubules (anti-Dlg1, green) form a network that extends throughout the myofibrils (phalloidin, F-actin, pink) in normal cardiomyocytes of the young adult dorsal vessel. Control, *w^1118^*. T-tubules were fragmented in cardiomyocytes of *Amph^Δ26^* mutants or with Hand4.2-GAL4 heart-targeted *PI4KIIIα* RNAi depletion. **(B)** Videography of live dorsal vessel imaged over 5 seconds in either control (*w1118)* or *Amph^Δ26^* mutant young eviscerated adult flies. **(C)** Heart rate (sec) obtained from high-speed imaging of the dorsal vessel (Fink et al., 2009) in 1 week-, 3 week- and 5 week-old flies for control Hand4.2-GAL4 and *Amph* RNAi conditions (left) and in control *w^1118^* and *Amph^Δ26^* mutant flies (right). All samples tested shown as scatter points in box and whisker plots with mean and Mann-Whitney test of significance.

Thus, both Drosophila body wall muscle and heart muscle require *PI4KIIIα* in a PI(4,5)P_2_ pathway for normal T-tubule membranes and muscle function. Our data in Drosophila together with previous work (Razzaq et al., 2001) suggests that PI(4,5)P_2_ and *Amph* have overlapping functions in growing muscle, as well as likely conserved mechanisms for T-tubule membrane organization shared through those important in human skeletal and heart muscles (Al-Qusairi and Laporte, 2011, Dibb et al., 2022).

The muscle similarities upon disruption of the *PI4KIIIα-sktl* pathway and *Amph* simply could be that PI(4,5)P_2_ is needed to recruit Amph and possibly other PIP-binding factors involved in muscle-specific membrane shaping and trafficking to form the T-tubule network (Lee et al., 2002, Vicinanza et al., 2008, Zhang and Zelhof, 2002). An alternative speculation could be that PI(4)P-PI(4,5)P_2_ synthesis serves directly as a critical gradient to drive lipid transport at the membrane contact sites (MCS) between the muscle plasma membrane and endoplasmic reticulum (Hammond and Burke, 2020). We showed that StmA specifically localized at the muscle dyad junctions along T-tubules. In this way, StmA recruitment and thus activation of PI4KIIIα at dyads could use phosphoinositide transport cycles to support changes in lipid composition for the vast expansion of the T-tubule membrane domains during larval muscle growth. In addition, our study revealed genetic distinctions between the identities of the longitudinal and transversal tubular membranes. Longitudinal and transversal elements have been noted to vary in timing with vertebrate muscle development and contexts of human muscle diseases (Al-Qusairi et al., 2009, Heinzel et al., 2008, Takekura et al., 2001, Wei et al., 2010, Wu et al., 2014a). Altogether, it remains to be tested whether the longitudinal tubular membranes serve as a lipid source or by other mechanisms as a precursor for the crucial transversal tubular membrane elements.

Here, we screened for T-tubule morphological differences to identify genes needed to form or maintain the larval muscle membrane network. Muscle-targeted RNAi depletion was able to recover specific T-tubule functions for genes that in null conditions were otherwise animal lethal (Yan et al., 2011), as well as determine a pathway that has relevance to muscle contraction and human myopathies (Al-Qusairi and Laporte, 2011, Dibb et al., 2022). Our previous work used a similar approach that identified gene functions specific for T-tubule remodeling at later developmental stages (Fujita et al., 2017). Altogether, the larval body wall muscles provide an experimental system useful to systematically interrogate, identify and distinguish genetic factors that shape, maintain or remodel the important T-tubule network.

## Materials and methods

### Drosophila strains and genetics

Flies were reared at 25°C, unless stated. Mef2-GAL4 driver was used for muscle-targeted gene expression. *UAS-LacZ* or *UAS-Luciferase* were used as a control in RNAi experiments. RNAi lines were obtained from Bloomington Drosophila Stock Center (BDSC, NIH P40OD018537) or Vienna Drosophila Resource Center (VDRC, www.vdrc.at), with the exception of *mtm* RNAi (Ribeiro et al., 2011) as listed on **Table 1**. Gene names used in the text and figures are as listed in FlyBase FB2023_06 (www.flybase.org) in order to conform with current standardized nomenclature. Note that RBO:eGFP was annotated here as StmA:GFP for consistency.

The following fly stocks were used: (1) *y[1] w[*]; P{w[+mC]=GAL4-Mef2.R}3*, referred to as *Mef2-GAL4* (BDSC_27390); (2) y[1] *w[*]; P{w[+mC]=UAS-mCD8::GFP.L}LL5, P{UAS-mCD8::GFP.L}2* (BDSC_5137); (3) *w[1118]; P{w[+mC]=UAS-Dcr-2.D}10* (BDSC_24651); (4) *UAS-eGFP*; (5) *P{PTT-GC}dlg1^YC0005^*, referred to as *Dlg1:GFP* (from L. Cooley, FlyTrap; (Quinones-Coello et al., 2007)); (6) *UASp-PLCδ-PH:GFP* (from J. Brill); (7) *UAS-mCherry:PLCδ-PH* (I. Ribeiro, A. Kiger); (8) *UAS-GFP:Tubby-Cter* (I. Ribeiro, A. Kiger); (9) *αTub84B-mCh:2xP4M* and *αTub84B-GFP:2xP4M* (J. Brill, (Ma et al., 2020)); (10) *UAS-YFP:PH-FAPP1* (I. Ribeiro, A. Kiger); (11) *UAS-GFP:PH-TAPP1* (I. Ribeiro, A. Kiger); (12) *UAS-GFP:myc:2xFYVE* (Wucherpfennig et al., 2003); (13) *UAS-PX-phox40:GFP* (I. Ribeiro, A. Kiger); (14) *w[1118]; tGPH^4^* (GFP:PH-Grp1; BDSC_8164); (15) *yw, rbo^2^; RBO:eGFP (T18),* under *rbo* promoter (from K. Broadie (Huang et al., 2004)); (16) *w; Rtnl:GFP^G00071^* (from L. Cooley, FlyTrap (Quinones-Coello et al., 2007); (17) *UAS-mCherry:jp*; (18) *UAS-GFP:jp*; (19) *w^[1118]^; PBac{y[+mDint2]w[+mC]=20XUAS-IVS-GCaMP6s}VK00005* (from J. Wang, BDSC_42749, (Chen et al., 2013)); (20) *w[1118]; Hand4.2-GAL4* (Han et al., 2006); (21) *y[1] v[1]; P{y[+t7.7]v[+t1.8]=UAS-LUC.VALIUM10}attP2* (UAS-Luciferase, BDSC_35788); (22) *UAS-IR-Amph* RNAi (*y[1] v[1]; P{TRiP.JF02883}attP2*, BDSC_28048); (23) *UAS-IR-Pi4KIIIα^v105614^* RNAi (VDRC_v105614); (24) *UAS-IR-Pi4KIIIα^v15993^* RNAi (VDRC_v15993) (25) *UAS-IR-Pi4KIIIα* RNAi (*y[1] sc[*] v[1] sev[21]; P{TRiP.HMS01686}attP40*; BDSC_38242); (26) *UAS-IR-sktl^v101624^* RNAi (VDRC_v101624); (27) *UAS-IR-sktl* RNAi (*y[1] v[1]; P{TRiP.JF02796}attP2,* BDSC_27715); (28) *UAS-IR-fwd* RNAi (*y[1] v[1]; P{TRiP.JF03329}attP2,* BDSC_29396); (29) *UAS-IR-PI4KIIα^v40995^* RNAi (VDRC_v40995); (30) *UAS-IR-PIP5K59^v47029^* RNAi (VDRC_v47029); (31) *UAS-IR-stmA* RNAi; (32) *UAS-IR-Ttc7* RNAi; (33) w*; Amph[26] (BDSC# 6498, (Razzaq et al., 2001)).

### RNAi screen

In primary screen, RNAi inverted repeat (IR) hairpins (**Table 1**) were crossed to *w, Dlg1:GFP; Mef2-GAL4, UAS-Dcr2* at 25°C. Live larvae were mounted for confocal microscopy imaging of Dlg1:GFP in muscle viewed through the cuticle. At least ten body wall muscles in three animals were checked for each genotype. T-tubule morphology was visually scored, noting differences compared to wildtype condition observed in tubule membrane pattern such as linear continuity, uniformity in size and spacing, and coverage across a muscle. Lethality was checked after two weeks from crossing. Phenotypes were validated by immunofluorescence microscopy of Dlg1 and F-actin in fixed and filleted larval muscle.

### Reagents and antibodies

Alexa Fluor 546 Phalloidin was used at 1.0 U/ml (Invitrogen). The following antibodies were used: mouse anti-fly Dlg1 (1:200; clone 4F3; Developmental Studies Hybridoma Bank), rabbit anti-Zormin B1 (Burkart et al., 2007) (1:500; B. Bullard, RRID:AB_2631283), rabbit anti-Amph Ra29 (Razzaq et al., 2001) (1:500; C. O’Kane), and secondary Alexa Fluor 546 or 647 antibodies (1:400; Molecular Probes).

### Muscle immunostaining

Third instar larvae were used at 5 days after egg laying (AEL), except when noted use of first or second instar larvae (30 and 60 hours AEL, Figure S1) or adult flies (1-week or 3-week-old, Figure S3). Preparations for imaging larval body wall muscles or adult dorsal vessels were performed as previously described (Fujita et al., 2017). Staged larvae were pinned on a sylgard-covered petri dish in dissection buffer (5 mM HEPES, 128 mM NaCl, 2 mM KCl, 4 mM MgCl_2_, 36 mM sucrose, pH 7.2). Larval bodies were opened with scissors, pinned flat, and fixed for 20 min. (4% formaldehyde, 50 mM EGTA, PBS). Then, the samples were unpinned and blocked for 30 min (0.3% bovine serum albumin (BSA), 2% goat serum, 0.6% Triton, PBS), incubated with primary antibody overnight at 4°C, washed (0.1% Triton PBS), then incubated for 2h at room temp with Alexa Fluor-conjugated secondary antibodies (1:400; Molecular Probes) and counterstained with phalloidin for F-actin (Invitrogen) as needed. The stained samples were washed and mounted in SlowFade reagent (Life Technologies; S36936) or FluorSave Reagent (Millipore; F345789).

### Confocal fluorescence microscopy

For imaging live larval body wall muscles, staged larvae were mounted using PBS between slide-glass and cover-glass following a protocol (Zitserman and Roegiers, 2011), and imaged from the dorsal side. Live or immunostained body wall muscles were acquired at room temperature (∼22°C) on a Zeiss LSM 700 laser confocal microscope with a 10x air/0.45 NA Plan Apochromatic objective or 40x oil/1.3 NA Plan Apochromatic objective. Image acquisition software used was Zen (Zeiss), and raw files were exported as 16 bit .tif files. The images were adjusted using Photoshop CS4 (Adobe) and Fiji ImageJ software.

### Electron microscopy

For TEM of larval body wall muscle, staged larvae were pinned on a sylgard-covered petri dish, dissected in fixative (2% paraformaldehyde, 2.5% glutaraldehyde, 150 mM sodium cacodylate, pH 7.4) and fixed for 1.5h at room temp. Then, samples were post-fixed in 1% osmium tetroxide in 0.1 M cacodylate buffer and stained in 1% uranyl acetate for 1h. Abdomen fillets were embedded in epoxy resin and 70 nm sections were collected on Formvar and carbon-coated copper grids. Images were acquired on a transmission electron microscope (FEI Tecnai Spirit G2 BioTWIN) and photographed by a bottom mount Eagle 4K digital camera.

### DNA engineering

Drosophila *junctophilin* isoform A (FBgn0032129, FlyBase) was PCR amplified from cDNA GH28348 (Drosophila Genomics Resource Center), cloned into pENTR/D-TOPO (LifeTechnologies), and subcloned by LR recombination into Gateway destination vectors to generate pUASt-GFP:jp and pUASt-mCherry:jp. Transgenic flies were generated following standard injection procedures (BestGene, Inc.).

### RT-PCR

Larvae were eviscerated to enrich for body wall muscle. Total RNA was isolated and reverse transcribed by random hexamers, and then cDNA was RT-PCR amplified and run on an agarose gel. RT-PCR product levels for *Amph*, *PI4KIIIα*, and *18S* rRNA were assessed from control larvae (*UAS-lacZ*) versus respective RNAi knockdown conditions. Primers used: *Amph* Forward #1532 5’-AACACGCTGGACGTGCCAAG-3’; *Amph*Reverse #1533 5’-CAATTTGTGCGCAAAGTCCGC-3’; *PI4KIIIα* Forward #1534 5’-CGCCTATCTGATCATATGCTCGG-3’; *PI4KIIIα* Reverse #1535 5’- AATGGGGCGCGATCAGTATGG-3’; *18S* Forward #1556 5’-GAAGTATGGTTGCAAAGCTGA-3’; *18S* Reverse #1557 5’-TGTTGTAAGTACTCGCCACA-3’.

### Larval roll-over assay

To assess larval muscle function, third instar larvae were washed with PBS and put on paper to remove PBS. Using blunt forceps, individual larvae were placed on 0.7% agar plate to dry for 1 min. Then, larvae were placed on another 0.7% agar plate upside down (with dorsal side down) onto an agarose plate, and timed how long it took to roll right-side up (with dorsal side up). Times were measured in seconds (s) and averaged for three attempts per larva with a minimum of 20 assays conducted per genotype.

### Calcium imaging and analysis

Staged third instar larvae expressing GCaMP6s (Chen et al., 2013) in muscle were mounted for live confocal microscopy imaging, as detailed above. Imaging was conducted for continuous seconds in each condition. Individual muscles undergoing spontaneous contraction were outlined to determine intensity of GCaMP6s within muscle area using ImageJ and plotted using Prism (GraphPad Software). The intensity of the highest frame (contracted muscle) was divided by the lowest frame (relaxed muscle), or ι1F/F0. 10 assays were conducted per genotype.

### In Situ Assessment of Cardiac Function

The cardiac activity of denervated Drosophila hearts was examined using an *in situ* dissection protocol to expose the autonomously beating heart (Fink et al., 2009, Ocorr et al., 2009, Vogler and Ocorr, 2009). For detailed cardiac parameter analysis, Semi-automated Optical Heartbeat Analysis (SOHA) was employed, enabling the evaluation of heart function via high-speed videographic recordings (Fink et al., 2009). The experimental procedure involved the transient immobilization of more than 15 *Drosophila* specimens using with 15µl FlyNap on filter paper. Subsequently, the flies were transferred to a 10mm x 35mm Petri dish, with application of a petroleum jelly to adhere the hydrophobic wing cuticle to the dish surface. To each dish, oxygenated artificial hemolymph at room temperature was added, composed of 108mM NaCl, 5mM KCl, 2mM CaCl_2_•2H_2_O, 8mM MgCl_2_•6H_2_O, 15mM pH 7.1 HEPES, 1mM NaH_2_PO_4_•H2O, 4mM NaHCO_3_, 10mM sucrose, and 5mM trehalose. Dissection procedures followed the protocol established by (Vogler and Ocorr, 2009), with a minimum 15-minute oxygenation period to ensure equilibrium. The dissected specimens were then recorded for 30 seconds using an Olympus BX63 microscope at 10X magnification, coupled with a Hamamatsu C9300 digital camera and HCImageLive software. Post-recording, the video files were processed through SOHA for manual identification of end diastolic and end systolic diameters at the ostia termini, subsequently facilitating the extraction of key cardiac function parameters (Fink et al., 2009).

### Statistics

Each experiment was performed at least three times as biological and technical replicates (at least three different cohorts of unique flies were analyzed in repeat procedures performed on at least three different days). One exception was for TEM analyses, which were performed on two parallel replicates with multiple animals each. All replicate experiments were performed in parallel with wildtype controls. *PI4KIIIα* RNAi phenotypes were penetrant. Data shown is representative and inclusive of all results for procedures with properly performing experimental controls. Statistical analysis was performed using a two-tailed unpaired *t* test in Prism (GraphPad Software).

Pearson’s correlation (*r*) measurements for co-localization quantification were performed on ImageJ software as shown in Figure 2B, 2H and Figure 3D. For each combination, ten 30 x 30 μm cropped images taken from different animals were used. Line scans in Figure 2D and 2E were performed on representative images for conditions with visually obvious colocalized fluorescence. Line scans were plotted as fluorescence intensity per channel at pixels along an arbitrary line manually drawn on images, as shown by yellow arrow on merged panel. Intensity graphed in arbitrary units (y) to overlay channels across pixels (x). Jp foci intensity (Figure 3F) and Jp overlap with Dlg1 T-tubules (Figure 3G) were analyzed by setting manual thresholds where the dimmest puncta were visible. Calcium flux analyzed as maximum change in GCaMP intensity plotted as scatter points with box and whisker plots and Mann-Whitney tests (Figure 4C). Heart function plotted as scatter points and assessed by Mann-Whitney test (Figure 5C) and Kruskal-Wallis test (Supplemental Figure S4A-S4F).

## Supplemental material

**Figure S1** shows specificity of the *PI4KIIIα* requirement and pathway in larval muscle T-tubule formation. Related to Figure 1. **Figure S2** provides a schematic of the seven phosphoinositides and possible interconversions by all predicted phosphoinositide kinases and phosphatases encoded in Drosophila, and the survey of different phosphoinositide biosensors in wildtype and mutant larval muscle. Related to Figure 2. **Figure S3** demonstrates a similar distribution of sarcoplasmic reticulum in wildtype and mutant muscle, and shows shared T-tubule phenotype of *Amph* RNAi in dorsal vessel to those from *Amph* null or *PI4KIIIα* RNAi. Related to Figure 3 and Figure 5. **Figure S4** shows results of heart function parameters measured from cardiac live imaging. Related to Figure 5. **Video 1** shows the confocal *z-*stack of Dlg1:GFP marking the T-tubule network in live dorsal body wall muscle imaged through the larval cuticle. Related to Figure 1. **Video 2** (control) and **Video 3** (*PI4KIIIα* RNAi) show views of calcium dynamics in larval dorsal body wall muscle detected by GCaMP intensity changes upon spontaneous muscle contractions. Related to Figure 4.

## Data availability statement

All associated data are available within the published report and online supplemental material. Any novel Drosophila strains generated in this study are available from the corresponding author upon reasonable request.

## Supporting information

Fujita bioRxiv Supplement Figures

Fujita bioRxiv Table 1

Fujita bioRxiv Video 1

Fujita bioRxiv Video 2

Fujita bioRxiv Video 3

## Acknowledgments

We are grateful to B. Bullard, J. Brill, K. Broadie, L. Cooley, M. Gonzalez-Gaitan, C. O’Kane, J. Wang, Bloomington Drosophila Stock Center, Drosophila Genomics Resource Center, Vienna Drosophila Stock Center, FlyTrap, Fly Stocks of National Institute of Genetics, and Developmental Studies Hybridoma Bank for reagents, and FlyBase for informational resources. We thank I. Ribeiro for generation of phosphoinositide biosensor transgenic lines while in the lab, and we thank T. Meerloo and M. Farquhar for technical assistance and use of the UCSD CMM Electron Microscope Facility. This work was supported in part by JSPS, the Uehara Memorial Foundation, and the Kanae Foundation postdoctoral fellowships to NF, and an American Heart Association grant 15IRG22830029 and National Institutes of Health grant R01AR073840 to AAK.

The authors declare no competing financial interests.

## Author contributions

Naonobu Fujita designed the research, performed the experiments, generated reagents, analyzed and interpreted data, and provided input on the manuscript.

Shravan Girada designed and performed experiments, analyzed and interpreted data, and provided input on the manuscript.

Georg Vogel performed experiments and analyzed and interpreted data.

Rolf Bodmer analyzed and interpreted data and provided input on the manuscript.

Amy A. Kiger designed the study, coordinated the project, analyzed and interpreted data, and wrote the manuscript.

## Abbreviations

PI(4)P: phosphatidylinositol 4-phosphate
PI(4,5)P_2_: phosphatidylinositol 4, 5-bisphosphate
PIP: phosphoinositide
SR: sarcoplasmic reticulum
T-tubule: Transverse tubule

## Figure legends

**Supplemental Figure S1. Knockdown of *PI4KIIIα-sktl* pathway has T-tubule-specific defects. (A)** RT-PCR of *Amph* and *PI4KIIIα* levels (left) and *18S* rRNA internal control (right) from larval body wall muscle of Control *UAS-lacZ* versus the respective RNAi knockdowns. **(B)** Top row: Control, *UAS-lacZ*. Larval dorsal body wall muscles at 1^st^, 2^nd^ and 3^rd^ instar stages (hours post-egg lay) detected by Mef2-Gal4-driven mCD8:GFP. Middle row: Control, *UAS-lacZ*. Dlg1:GFP-positive T-tubule network with transversal and longitudinal elements at all stages. Bottom row: Mef2-Gal4-driven *PI4KIIIα* RNAi. Disrupted larval Dlg1:GFP-marked T-tubule network with few transversal elements by 3^rd^ instar stage. **(C)** T-tubule organization (anti-Dlg) was unaffected by RNAi knockdown of *fwd* (PI4KIIIβ), *PI4KII* or *PIP5K59*. **(D)** Organization of T-tubules (anti-Dlg, green) and F-actin (phalloidin, pink) in fixed muscles from control or knockdown conditions: *Amph* RNAi, *PI4KIIIα* RNAi, *stmA* RNAi, *Ttc7* RNAi or *sktl* RNAi. All conditions contained myofibrils. Scale bars 10 µm, except as indicated 500 µm in (B).

**Supplemental Figure S2. Survey identified PI(4)P and PI(4,5)P_2_ at T-tubules dependent on PI4KIIIα pathway. (A)** Seven phosphoinositides, the phosphorylated forms of PtdIns, are interconverted by families of specific PI-kinases and PI-phosphatases, as shown for the enzyme activities encoded in Drosophila. **(B)** Mef2-GAL4 larval muscle expression of UAS-driven phosphoinositide biosensors surveyed for T-tubule localization of PI(3)P [GFP:2xFYVE and PX-phox40:GFP], PI(4)P [YFP:PH-FAPP1 and GFP:P4M], PI(3,4)P_2_ [GFP:TAPP1-PH], PI(4,5)P_2_ [GFP:Tubby-C-PH and GFP:PH-PLC8], and PI(3,4,5)P_3_ [PH-Grp1:GFP]. Only P4M and both PI(4,5)P_2_ biosensors were detected at T-tubules. **(C)** PI(4,5)P_2_ detected by mCherry:PH-PLC8 was present at T-tubule membranes in control larval muscle (top) and absent with *stmA* depletion (middle). Although T-tubule network was disrupted with *Amph* RNAi, PI(4,5)P_2_ detected by mCherry:PH-PLC8 was still present at longitudinal T-tubule membrane remnants (bottom). Scale bars 20 µm.

**Supplemental Figure S3. Muscle with RNAi loss of transversal T-tubule membranes maintain sarcoplasmic reticulum organization. (A)** Merge and dual channel overlays for same data shown in Figure 3A: Junctophilin (mCherry:Jp, pink) at dyad junctions that form between the sarcoplasmic reticulum (SR; Rtnl1:GFP, green) and T-tubule membranes (anti-Dlg1, cyan) along both transversal (yellow arrowheads) and longitudinal (white arrowheads) elements. **(B)** SR marked by Rtnl1:GFP (green) distributed between and alongside the intact T-tubule network (anti-Dlg1, pink) in control muscle (top row); similar overall SR distribution seen in *PI4KIIIα* RNAi or *Amph* RNAi muscles despite fragmented longitudinal T-tubules (middle and lower rows, respectively). White arrowheads: examples of Rtnl:GFP and Dlg1 longitudinal membrane contact sites. Yellow arrowheads: examples of Rtnl:GFP between longitudinal membranes. **(C)** Individual channel and merged images of T-tubules (anti-Dlg1, green) and myofibrils (phalloidin, pink) in cardiomyocytes of adult dorsal vessel. Control (*Hand-GAL4 UAS-Luciferase*), top: T-tubules form a continuous network in cardiomyocytes of 1-week and 3-week old adult control flies. *Amph-*depleted cardiomyocytes (*Hand-GAL4 UAS-Amph* RNAi), bottom: fragmented T-tubules in cardiomyocytes of 1-week and 3-week old adult flies, similar to *Amph^Δ26^* genomic mutant flies (shown in Figure 5A). Scale bars 5 µm (A, B) and 10 µm (C).

**Supplemental Figure S5. Dorsal vessels with disrupted T-tubules and decreased heart rate maintain overall cardiomyocyte morphology and contractability.** High-speed imaging from 1 week- and 3 week-old flies analyzed for parameters of heart function in control Hand4.2-GAL4 and *Amph* RNAi conditions (left), and in control *w^1118^* and *Amph^Δ26^* mutant flies (right). All samples tested shown as scatter plots with Kruskal-Wallis test of significance. Besides decreased heart rate (Figure 5C), the only differences seen in heart function were increased heart period, diastolic intervals and systolic intervals in 1 week-old *Amph^Δ26^* mutant flies. **(A)** Heart period mean (sec). **(B)** Diastolic intervals (sec). **(C)** Systolic intervals (sec). **(D)** Fractional shortening. **(E)** Diastolic diameter (µm). **(F)** Systolic diameter (µm). ns = not significant.

**Supplemental video 1. Image *z*-stack of Dlg1:GFP labelled T-tubules in live larval dorsal muscle.** Confocal microscopy *z-*stack (10 x 2 µm) of Dlg1:GFP indicating the vast, elaborate T-tubule network in live dorsal body wall muscle imaged through the third instar larval cuticle. Related to Figure 1A.

**Supplemental video 2. Calcium dynamics (GCaMP) with control larval muscle contractions.** Confocal microscopy of control live third instar larva with increases in fluorescent calcium reporter, GCaMP, indicating cytoplasmic calcium flux with body wall muscle contractions. Related to Figure 4.

**Supplemental video 3. Calcium dynamics (GCaMP) in *PI4KIIIα* RNAi larval muscles.** Confocal microscopy of live third instar larva with muscle-targeted Mef2-GAL4 *PI4KIIIα* RNAi, showing less intense pulses of fluorescent calcium reporter, GCaMP, with weaker body wall muscle contractions than in control animals. Related to Figure 4.

## References

Al-Qusairi, L. & Laporte, J. 2011. T-tubule biogenesis and triad formation in skeletal muscle and implication in human diseases. Skeletal Muscle, 1, 26.

Al-Qusairi, L., Weiss, N., Toussaint, A., Berbey, C., Messaddeq, N., Kretz, C., Sanoudou, D., Beggs, A. H., Allard, B., Mandel, J. L., Laporte, J., Jacquemond, V. & Buj-Bello, A. 2009. T-tubule disorganization and defective excitation-contraction coupling in muscle fibers lacking myotubularin lipid phosphatase. Proc Natl Acad Sci U S A, 106, 18763–8.

Baird, D., Stefan, C., Audhya, A., Weys, S. & Emr, S. D. 2008. Assembly of the PtdIns 4-kinase Stt4 complex at the plasma membrane requires Ypp1 and Efr3. J Cell Biol, 183, 1061–74.

Balakrishnan, S. S., Basu, U., Shinde, D., Thakur, R., Jaiswal, M. & Raghu, P. 2018. Regulation of PI4P levels by PI4KIIIalpha during G-protein-coupled PLC signaling in Drosophila photoreceptors. J Cell Sci, 131.

Balla, T. 2013. Phosphoinositides: tiny lipids with giant impact on cell regulation. Physiol Rev, 93, 1019–137.

Bashir, R., Britton, S., Strachan, T., Keers, S., Vafiadaki, E., Lako, M., Richard, I., Marchand, S., Bourg, N., Argov, Z., Sadeh, M., Mahjneh, I., Marconi, G., Passos-Bueno, M. R., Moreira Ede, S., Zatz, M., Beckmann, J. S. & Bushby, K. 1998. A gene related to Caenorhabditis elegans spermatogenesis factor fer-1 is mutated in limb-girdle muscular dystrophy type 2B. Nat Genet, 20, 37–42.

Betz, R. C., Schoser, B. G., Kasper, D., Ricker, K., Ramirez, A., Stein, V., Torbergsen, T., Lee, Y. A., Nothen, M. M., Wienker, T. F., Malin, J. P., Propping, P., Reis, A., Mortier, W., Jentsch, T. J., Vorgerd, M. & Kubisch, C. 2001. Mutations in CAV3 cause mechanical hyperirritability of skeletal muscle in rippling muscle disease. Nat Genet, 28, 218–9.

Bitoun, M., Maugenre, S., Jeannet, P. Y., Lacene, E., Ferrer, X., Laforet, P., Martin, J. J., Laporte, J., Lochmuller, H., Beggs, A. H., Fardeau, M., Eymard, B., Romero, N. B. & Guicheney, P. 2005. Mutations in dynamin 2 cause dominant centronuclear myopathy. Nat Genet, 37, 1207–9.

Blondeau, F., Laporte, J., Bodin, S., Superti-Furga, G., Payrastre, B. & Mandel, J. L. 2000. Myotubularin, a phosphatase deficient in myotubular myopathy, acts on phosphatidylinositol 3-kinase and phosphatidylinositol 3-phosphate pathway. Hum Mol Genet, 9, 2223–9.

Brette, F. & Orchard, C. 2007. Resurgence of cardiac t-tubule research. Physiology (Bethesda), 22, 167–73.

Brombacher, E., Urwyler, S., Ragaz, C., Weber, S. S., Kami, K., Overduin, M. & Hilbi, H. 2009. Rab1 guanine nucleotide exchange factor SidM is a major phosphatidylinositol 4-phosphate-binding effector protein of Legionella pneumophila. J Biol Chem, 284, 4846–56.

Burkart, C., Qiu, F., Brendel, S., Benes, V., Haag, P., Labeit, S., Leonard, K. & Bullard, B. 2007. Modular proteins from the Drosophila sallimus (sls) gene and their expression in muscles with different extensibility. J Mol Biol, 367, 953–69.

Calpena, E., Lopez Del Amo, V., Chakraborty, M., Llamusi, B., Artero, R., Espinos, C. & Galindo, M. I. 2018. The Drosophila junctophilin gene is functionally equivalent to its four mammalian counterparts and is a modifier of a Huntingtin poly-Q expansion and the Notch pathway. Dis Model Mech, 11.

Chen, T. W., Wardill, T. J., Sun, Y., Pulver, S. R., Renninger, S. L., Baohan, A., Schreiter, E. R., Kerr, R. A., Orger, M. B., Jayaraman, V., Looger, L. L., Svoboda, K. & Kim, D. S. 2013. Ultrasensitive fluorescent proteins for imaging neuronal activity. Nature, 499, 295–300.

Chin, Y. H., Lee, A., Kan, H. W., Laiman, J., Chuang, M. C., Hsieh, S. T. & Liu, Y. W. 2015. Dynamin-2 mutations associated with centronuclear myopathy are hypermorphic and lead to T-tubule fragmentation. Hum Mol Genet, 24, 5542–54.

Devergne, O., Tsung, K., Barcelo, G. & Schupbach, T. 2014. Polarized deposition of basement membrane proteins depends on Phosphatidylinositol synthase and the levels of Phosphatidylinositol 4,5-bisphosphate. Proc Natl Acad Sci U S A, 111, 7689–94.

Dibb, K. M., Louch, W. E. & Trafford, A. W. 2022. Cardiac Transverse Tubules in Physiology and Heart Failure. Annu Rev Physiol, 84, 229–255.

Dowling, J. J., Vreede, A. P., Low, S. E., Gibbs, E. M., Kuwada, J. Y., Bonnemann, C. G. & Feldman, E. L. 2009. Loss of myotubularin function results in T-tubule disorganization in zebrafish and human myotubular myopathy. PLoS Genet, 5, e1000372.

Fink, M., Callol-Massot, C., Chu, A., Ruiz-Lozano, P., Izpisua Belmonte, J. C., Giles, W., Bodmer, R. & Ocorr, K. 2009. A new method for detection and quantification of heartbeat parameters in Drosophila, zebrafish, and embryonic mouse hearts. Biotechniques, 46, 101–13.

Fugier, C., Klein, A. F., Hammer, C., Vassilopoulos, S., Ivarsson, Y., Toussaint, A., Tosch, V., Vignaud, A., Ferry, A., Messaddeq, N., Kokunai, Y., Tsuburaya, R., De La Grange, P., Dembele, D., Francois, V., Precigout, G., Boulade-Ladame, C., Hummel, M. C., Lopez De Munain, A., Sergeant, N., Laquerriere, A., Thibault, C., Deryckere, F., Auboeuf, D., Garcia, L., Zimmermann, P., Udd, B., Schoser, B., Takahashi, M. P., Nishino, I., Bassez, G., Laporte, J., Furling, D. & Charlet-Berguerand, N. 2011. Misregulated alternative splicing of BIN1 is associated with T tubule alterations and muscle weakness in myotonic dystrophy. Nat Med, 17, 720–5.

Fujita, N., Huang, W., Lin, T. H., Groulx, J. F., Jean, S., Nguyen, J., Kuchitsu, Y., Koyama-Honda, I., Mizushima, N., Fukuda, M. & Kiger, A. A. 2017. Genetic screen in Drosophila muscle identifies autophagy-mediated T-tubule remodeling and a Rab2 role in autophagy. Elife, 6.

Galbiati, F., Engelman, J. A., Volonte, D., Zhang, X. L., Minetti, C., Li, M., Hou, H., JR., Kneitz, B., Edelmann, W. & Lisanti, M. P. 2001. Caveolin-3 null mice show a loss of caveolae, changes in the microdomain distribution of the dystrophin-glycoprotein complex, and t-tubule abnormalities. J Biol Chem, 276, 21425–33.

Hall, T. E., Martel, N., Ariotti, N., Xiong, Z., Lo, H. P., Ferguson, C., Rae, J., Lim, Y. W. & Parton, R. G. 2020. In vivo cell biological screening identifies an endocytic capture mechanism for T-tubule formation. Nat Commun, 11, 3711.

Hammond, G. R., Machner, M. P. & Balla, T. 2014. A novel probe for phosphatidylinositol 4-phosphate reveals multiple pools beyond the Golgi. J Cell Biol, 205, 113–26.

Hammond, G. R. V. & Burke, J. E. 2020. Novel roles of phosphoinositides in signaling, lipid transport, and disease. Curr Opin Cell Biol, 63, 57–67.

Han, Z., Yi, P., Li, X. & Olson, E. N. 2006. Hand, an evolutionarily conserved bHLH transcription factor required for Drosophila cardiogenesis and hematopoiesis. Development, 133, 1175–82.

Heinzel, F. R., Bito, V., Biesmans, L., Wu, M., Detre, E., Von Wegner, F., Claus, P., Dymarkowski, S., Maes, F., Bogaert, J., Rademakers, F., D’hooge, J. & Sipido, K. 2008. Remodeling of T-tubules and reduced synchrony of Ca2+ release in myocytes from chronically ischemic myocardium. Circ Res, 102, 338–46.

Hofhuis, J., Bersch, K., Bussenschutt, R., Drzymalski, M., Liebetanz, D., Nikolaev, V. O., Wagner, S., Maier, L. S., Gartner, J., Klinge, L. & Thoms, S. 2017. Dysferlin mediates membrane tubulation and links T-tubule biogenesis to muscular dystrophy. J Cell Sci, 130, 841–852.

Huang, F. D., Matthies, H. J., Speese, S. D., Smith, M. A. & Broadie, K. 2004. Rolling blackout, a newly identified PIP2-DAG pathway lipase required for Drosophila phototransduction. Nat Neurosci, 7, 1070–8.

Jensen, H. 1977. Ultrastructure of the myocardial cell and its membrane systems in the adult fly Calliphora erythrocephala (insecta: diptera). Cell Tissue Res, 180, 293–302.

Katan, M. & Cockcroft, S. 2020. Phosphatidylinositol(4,5)bisphosphate: diverse functions at the plasma membrane. Essays Biochem, 64, 513–531.

Klinge, L., Harris, J., Sewry, C., Charlton, R., Anderson, L., Laval, S., Chiu, Y. H., Hornsey, M., Straub, V., Barresi, R., Lochmuller, H. & Bushby, K.i 2010. Dysferlin associates with the developing T-tubule system in rodent and human skeletal muscle. Muscle Nerve, 41, 166–73.

Kwok, E., Otto, S. C., Khuu, P., Carpenter, A. P., Codding, S. J., Reardon, P. N., Vanegas, J., Kumar, T. M., Kuykendall, C. J., Mehl, R. A., Baio, J. & Johnson, C. P. 2023. The Dysferlin C2A Domain Binds PI(4,5)P2 and Penetrates Membranes. J Mol Biol, 435, 168193.

Laporte, J., Bedez, F., Bolino, A. & Mandel, J. L. 2003. Myotubularins, a large disease-associated family of cooperating catalytically active and inactive phosphoinositides phosphatases. Hum Mol Genet, 12 Spec No 2, R285–92.

Laporte, J., Hu, L. J., Kretz, C., Mandel, J. L., Kioschis, P., Coy, J. F., Klauck, S. M., Poustka, A. & Dahl, N. 1996. A gene mutated in X-linked myotubular myopathy defines a new putative tyrosine phosphatase family conserved in yeast. Nat Genet, 13, 175–82.

Lee, E., Marcucci, M., Daniell, L., Pypaert, M., Weisz, O. A., Ochoa, G. C., Farsad, K., Wenk, M. R. & De Camilli, P. 2002. Amphiphysin 2 (Bin1) and T-tubule biogenesis in muscle. Science, 297, 1193–6.

Lehmacher, C., Abeln, B. & Paululat, A. 2012. The ultrastructure of Drosophila heart cells. Arthropod Struct Dev, 41, 459–74.

Lipsett, D. B., Frisk, M., Aronsen, J. M., Norden, E. S., Buonarati, O. R., Cataliotti, A., Hell, J. W., Sjaastad, I., Christensen, G. & Louch, W. E. 2019. Cardiomyocyte substructure reverts to an immature phenotype during heart failure. J Physiol, 597, 1833–1853.

Liu, C. H., Bollepalli, M. K., Long, S. V., Asteriti, S., Tan, J., Brill, J. A. & Hardie, R. C. 2018. Genetic dissection of the phosphoinositide cycle in Drosophila photoreceptors. J Cell Sci, 131.

Liu, J., Aoki, M., Illa, I., Wu, C., Fardeau, M., Angelini, C., Serrano, C., Urtizberea, J. A., Hentati, F., Hamida, M. B., Bohlega, S., Culper, E. J., Amato, A. A., Bossie, K., Oeltjen, J., Bejaoui, K., Mckenna-Yasek, D., Hosler, B. A., Schurr, E., Arahata, K., De Jong, P. J. & Brown, R. H., JR. 1998. Dysferlin, a novel skeletal muscle gene, is mutated in Miyoshi myopathy and limb girdle muscular dystrophy. Nat Genet, 20, 31–6.

Ma, C. J., Yang, Y., Kim, T., Chen, C. H., Polevoy, G., Vissa, M., Burgess, J. & Brill, J. A. 2020. An early endosome-derived retrograde trafficking pathway promotes secretory granule maturation. J Cell Biol, 219.

Minetti, C., Sotgia, F., Bruno, C., Scartezzini, P., Broda, P., Bado, M., Masetti, E., Mazzocco, M., Egeo, A., Donati, M. A., Volonte, D., Galbiati, F., Cordone, G., Bricarelli, F. D., Lisanti, M. P. & Zara, F. 1998. Mutations in the caveolin-3 gene cause autosomal dominant limb-girdle muscular dystrophy. Nat Genet, 18, 365–8.

Muller, A. J., Baker, J. F., Duhadaway, J. B., Ge, K., Farmer, G., Donover, P. S., Meade, R., Reid, C., Grzanna, R., Roach, A. H., Shah, N., Soler, A. P. & Prendergast, G. C. 2003. Targeted disruption of the murine Bin1/Amphiphysin II gene does not disable endocytosis but results in embryonic cardiomyopathy with aberrant myofibril formation. Mol Cell Biol, 23, 4295–306.

Nakai, J., Ohkura, M. & Imoto, K. 2001. A high signal-to-noise Ca(2+) probe composed of a single green fluorescent protein. Nat Biotechnol, 19, 137–41.

Nakatsu, F., Baskin, J. M., Chung, J., Tanner, L. B., Shui, G., Lee, S. Y., Pirruccello, M., Hao, M., Ingolia, N. T., Wenk, M. R. & De Camilli, P. 2012. PtdIns4P synthesis by PI4KIIIalpha at the plasma membrane and its impact on plasma membrane identity. J Cell Biol, 199, 1003–16.

Nicot, A. S., Toussaint, A., Tosch, V., Kretz, C., Wallgren-Pettersson, C., Iwarsson, E., Kingston, H., Garnier, J. M., Biancalana, V., Oldfors, A., Mandel, J. L. & Laporte, J. 2007. Mutations in amphiphysin 2 (BIN1) disrupt interaction with dynamin 2 and cause autosomal recessive centronuclear myopathy. Nat Genet, 39, 1134–9.

Ocorr, K., Fink, M., Cammarato, A., Bernstein, S. & Bodmer, R. 2009. Semi-automated Optical Heartbeat Analysis of small hearts. J Vis Exp.

Peachey, L. D. 1968. Muscle. Annu Rev Physiol, 30, 401–40.

Piggott, C. A. & Jin, Y. 2021. Junctophilins: Key Membrane Tethers in Muscles and Neurons. Front Mol Neurosci, 14, 709390.

Posey, A. D., JR., Swanson, K. E., Alvarez, M. G., Krishnan, S., Earley, J. U., Band, H., Pytel, P., Mcnally, E. M. & Demonbreun, A. R. 2014. EHD1 mediates vesicle trafficking required for normal muscle growth and transverse tubule development. Dev Biol, 387, 179–90.

Quinones-Coello, A. T., Petrella, L. N., Ayers, K., Melillo, A., Mazzalupo, S., Hudson, A. M., Wang, S., Castiblanco, C., Buszczak, M., Hoskins, R. A. & Cooley, L. 2007. Exploring strategies for protein trapping in Drosophila. Genetics, 175, 1089–104.

Razzaq, A., Robinson, I. M., Mcmahon, H. T., Skepper, J. N., Su, Y., Zelhof, A. C., Jackson, A. P., Gay, N. J. & O’kane, C. J. 2001. Amphiphysin is necessary for organization of the excitation-contraction coupling machinery of muscles, but not for synaptic vesicle endocytosis in Drosophila. Genes Dev, 15, 2967–79.

Ribeiro, I., Yuan, L., Tanentzapf, G., Dowling, J. J. & Kiger, A. 2011. Phosphoinositide regulation of integrin trafficking required for muscle attachment and maintenance. PLoS Genet, 7, e1001295.

Rosemblatt, M., Hidalgo, C., Vergara, C. & Ikemoto, N. 1981. Immunological and biochemical properties of transverse tubule membranes isolated from rabbit skeletal muscle. J Biol Chem, 256, 8140–8.

Santagata, S., Boggon, T. J., Baird, C. L., Gomez, C. A., Zhao, J., Shan, W. S., Myszka, D. G. & Shapiro, L. 2001. G-protein signaling through tubby proteins. Science, 292, 2041–50.

Smith, D. S. 1966. The organization and function of the sarcoplasmic reticulum and T-system of muscle cells. Prog Biophys Mol Biol, 16, 107–42.

Souidi, A. & Jagla, K. 2021. Drosophila Heart as a Model for Cardiac Development and Diseases. Cells, 10.

Stauffer, T. P., Ahn, S. & Meyer, T. 1998. Receptor-induced transient reduction in plasma membrane PtdIns(4,5)P2 concentration monitored in living cells. Curr Biol, 8, 343–6.

Szentesi, P., Dienes, B., Kutchukian, C., Czirjak, T., Buj-Bello, A., Jacquemond, V. & Csernoch, L. 2023. Disrupted T-tubular network accounts for asynchronous calcium release in MTM1-deficient skeletal muscle. J Physiol, 601, 99–121.

Takekura, H., Flucher, B. E. & Franzini-Armstrong, C. 2001. Sequential docking, molecular differentiation, and positioning of T-Tubule/SR junctions in developing mouse skeletal muscle. Dev Biol, 239, 204–14.

Takekura, H. & Franzini-Armstrong, C. 2002. The structure of Ca(2+) release units in arthropod body muscle indicates an indirect mechanism for excitation-contraction coupling. Biophys J, 83, 2742–53.

Takeshima, H., Komazaki, S., Nishi, M., Iino, M. & Kangawa, K. 2000. Junctophilins: a novel family of junctional membrane complex proteins. Mol Cell, 6, 11–22.

Tan, J., Oh, K., Burgess, J., Hipfner, D. R. & Brill, J. A. 2014. PI4KIIIalpha is required for cortical integrity and cell polarity during Drosophila oogenesis. J Cell Sci, 127, 954–66.

Taylor, G. S., Maehama, T. & Dixon, J. E. 2000. Inaugural article: myotubularin, a protein tyrosine phosphatase mutated in myotubular myopathy, dephosphorylates the lipid second messenger, phosphatidylinositol 3-phosphate. Proc Natl Acad Sci U S A, 97, 8910–5.

Thallmair, V., Schultz, L., Zhao, W., Marrink, S. J., Oliver, D. & Thallmair, S. 2022. Two cooperative binding sites sensitize PI(4,5)P(2) recognition by the tubby domain. Sci Adv, 8, eabp9471.

Tian, L., Hires, S. A., Mao, T., Huber, D., Chiappe, M. E., Chalasani, S. H., Petreanu, L., Akerboom, J., Mckinney, S. A., Schreiter, E. R., Bargmann, C. I., Jayaraman, V., Svoboda, K. & Looger, L. L. 2009. Imaging neural activity in worms, flies and mice with improved GCaMP calcium indicators. Nat Methods, 6, 875–81.

Varnai, P. & Balla, T. 1998. Visualization of phosphoinositides that bind pleckstrin homology domains: calcium- and agonist-induced dynamic changes and relationship to myo-[3H]inositol-labeled phosphoinositide pools. J Cell Biol, 143, 501–10.

Vicinanza, M., D’angelo, G., Di Campli, A. & De Matteis, M. A. 2008. Function and dysfunction of the PI system in membrane trafficking. EMBO J, 27, 2457–70.

Voelker, T. L., Del Villar, S. G., Westhoff, M., Costa, A. D., Coleman, A. M., Hell, J. W., Horne, M. C., Dickson, E. J. & Dixon, R. E. 2023. Acute phosphatidylinositol 4,5 bisphosphate depletion destabilizes sarcolemmal expression of cardiac L-type Ca(2+) channel Ca(V)1.2. Proc Natl Acad Sci U S A, 120, e2221242120.

Vogler, G. & Ocorr, K. 2009. Visualizing the beating heart in Drosophila. J Vis Exp.

Wei, S., Guo, A., Chen, B., Kutschke, W., Xie, Y. P., Zimmerman, K., Weiss, R. M., Anderson, M. E., Cheng, H. & Song, L. S. 2010. T-tubule remodeling during transition from hypertrophy to heart failure. Circ Res, 107, 520–31.

Wu, C. Y., Chen, B., Jiang, Y. P., Jia, Z., Martin, D. W., Liu, S., Entcheva, E., Song, L. S. & Lin, R. Z. 2014a. Calpain-dependent cleavage of junctophilin-2 and T-tubule remodeling in a mouse model of reversible heart failure. J Am Heart Assoc, 3, e000527.

Wu, X., Chi, R. J., Baskin, J. M., Lucast, L., Burd, C. G., De Camilli, P. & Reinisch, K. M. 2014b. Structural insights into assembly and regulation of the plasma membrane phosphatidylinositol 4-kinase complex. Dev Cell, 28, 19–29.

Wucherpfennig, T., Wilsch-Brauninger, M. & Gonzalez-Gaitan, M. 2003. Role of Drosophila Rab5 during endosomal trafficking at the synapse and evoked neurotransmitter release. J Cell Biol, 161, 609–24.

Yan, Y., Denef, N., Tang, C. & Schüpbach, T. 2011. Drosophila PI4KIIIalpha is required in follicle cells for oocyte polarization and Hippo signaling. Development, 138, 1697–1703.

Zhang, B. & Zelhof, A. C. 2002. Amphiphysins: raising the BAR for synaptic vesicle recycling and membrane dynamics. Bin-Amphiphysin-Rvsp. Traffic, 3, 452–60.

Zhang, X., Wang, W. A., Jiang, L. X., Liu, H. Y., Zhang, B. Z., Lim, N., Li, Q. Y. & Huang, F. D. 2017. Downregulation of RBO-PI4KIIIalpha Facilitates Abeta(42) Secretion and Ameliorates Neural Deficits in Abeta(42)-Expressing Drosophila. J Neurosci, 37, 4928–4941.

Zitserman, D. & Roegiers, F. 2011. Live-cell imaging of sensory organ precursor cells in intact Drosophila pupae. J Vis Exp.

