## Supplementary figures and images for "PI(4,5)P_2_ role in Transverse-tubule membrane formation and muscle function"

### Fujita bioRxiv Supplement Figures

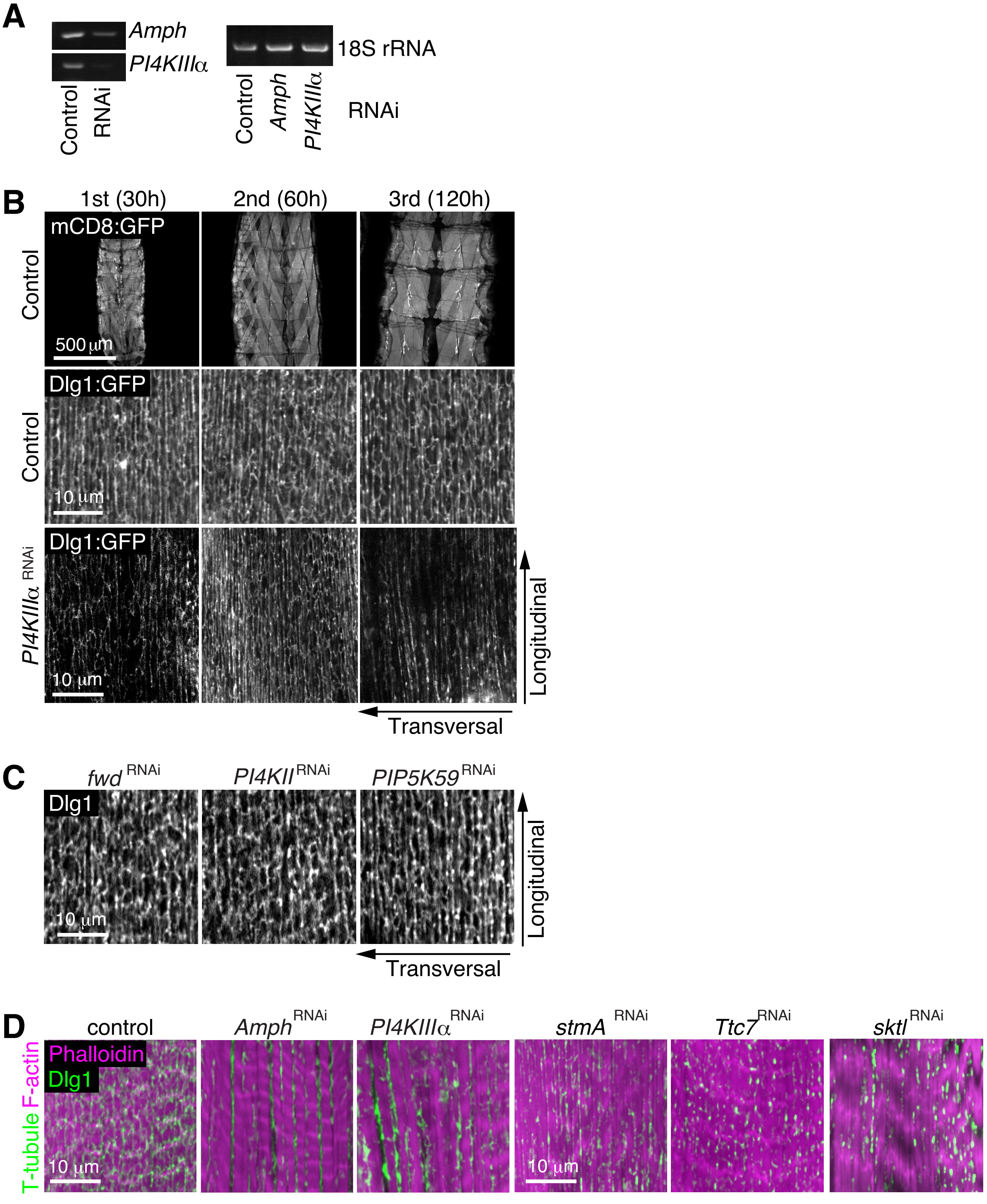

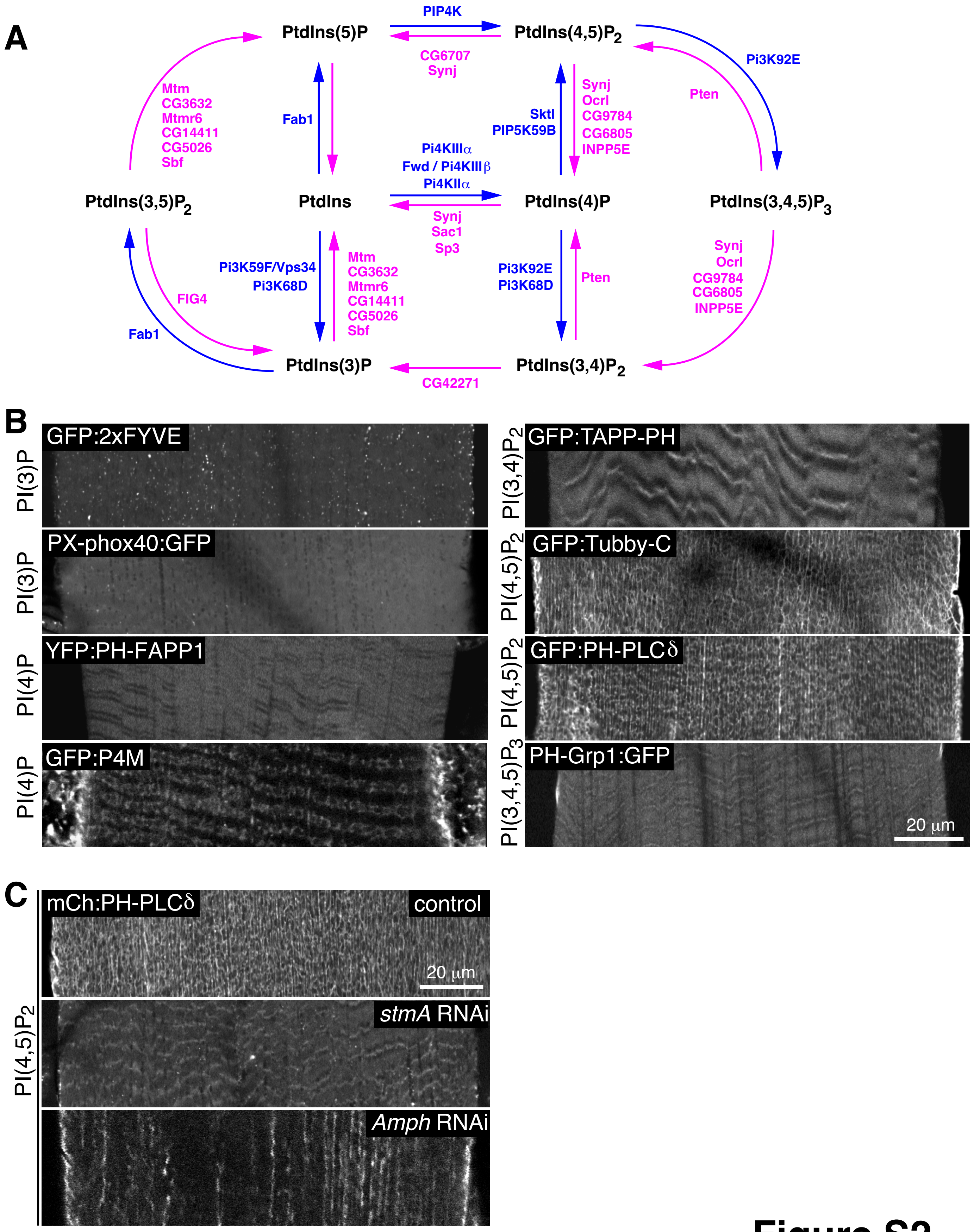

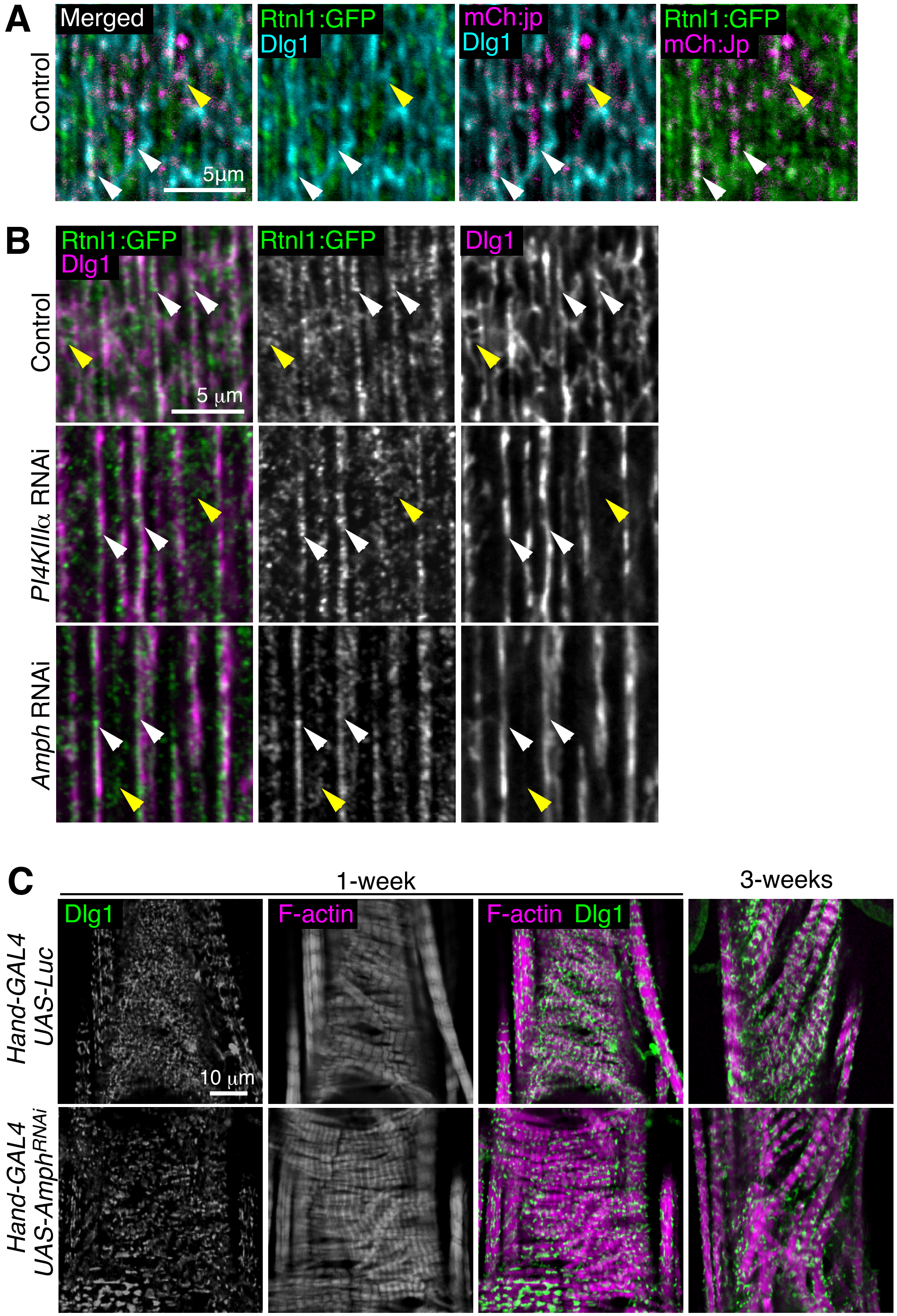

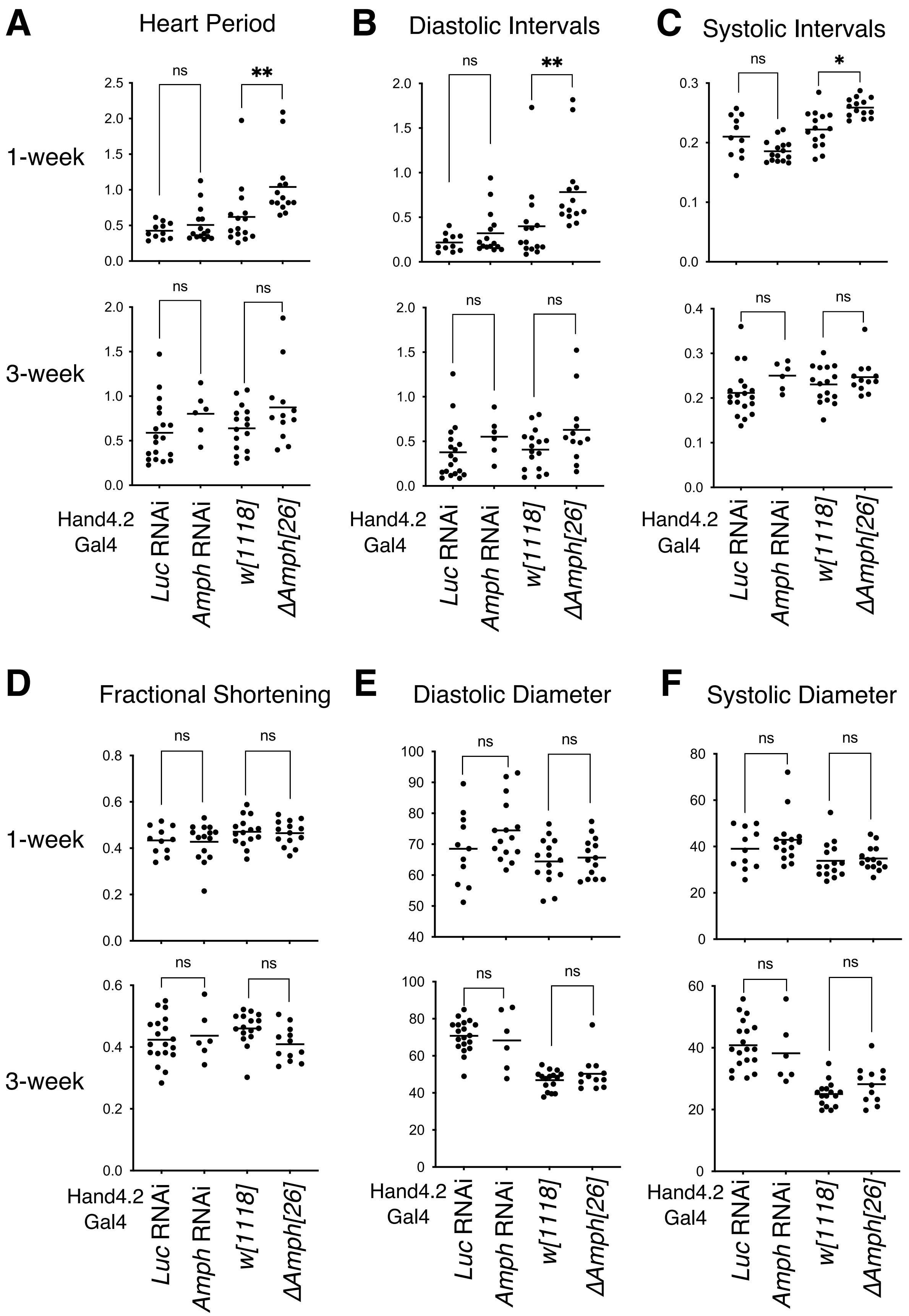
